# Redundant prefrontal hemispheres adapt storage strategy to working memory demands

**DOI:** 10.1101/2025.01.15.633176

**Authors:** Melanie Tschiersch, Akash Umakantha, Ryan C. Williamson, Matthew A. Smith, Joao Barbosa, Albert Compte

**Affiliations:** IDIBAPS, Barcelona, Spain; Programa de doctorat en Biomedicina, Universitat de Barcelona (UB), Barcelona, Spain; Center for the Neural Basis of Cognition, Carnegie Mellon University & University of Pittsburgh, Pittsburgh PA, USA; Carnegie Mellon University Neuroscience Institute, Pittsburgh PA, USA; Carnegie Mellon University Department of Biomedical Engineering, Pittsburgh PA, USA; Laboratoire de Neurosciences Cognitives et Computationnelles, INSERM U960, Ecole Normale Superieure - PSL Research University, 75005, Paris, France; Cognitive Neuroimaging Unit, INSERM, CEA, CNRS, Université Paris-Saclay, NeuroSpin center, 91191 Gif/Yvette, France; Institut de neuromodulation, GHU Paris, psychiatrie et neurosciences, centre hospitalier Sainte-Anne, pôle hospitalo-universitaire 15, Université Paris Cité, Paris, France; Institut d’Investigacions Biomèdiques de Barcelona (IIBB), CSIC, Barcelona, Spain

**Keywords:** attractor networks, working memory, prefrontal cortex, serial dependence, bilateral field advantage, multi-area recordings

## Abstract

The prefrontal hemispheres must coordinate dynamically to maintain a unified representation of visual space. Recently, two opposing theories using distinct storage strategies have been proposed: A high-capacity specialized architecture, where each hemisphere governs contralateral behavior, and a fail-safe redundant one, where both hemispheres jointly guide behavior across the visual space. Here, we analyzed simultaneous bilateral prefrontal cortex recordings from three macaque monkeys performing a visuo-spatial working memory task. Both hemispheres equally predicted behavioral imprecision, decoding errors were weakly correlated between hemispheres, and serial dependence remained local within hemispheres, suggesting a redundant, weakly coupled organization. Attractor network simulations showed that redundancy improved precision when task demands were below memory capacity, while weak interhemispheric coupling increased capacity in more demanding tasks by allowing hemispheric specialization. These predicted patterns were validated in human and monkey data, reconciling previous findings and revealing a versatile interhemispheric architecture that adapts to varying cognitive demands.

## Introduction

The world is spatially continuous, yet the inputs to the primary visual cortex (V1) are divided at the vertical meridian, with each hemisphere of the brain processing only the contralateral visual field (1). These divided hemifield representations are stitched together by callosal projections between the V1 hemispheres, which selectively connect neurons with receptive fields along the vertical midline (2–4). While this division is well understood in early visual areas, it remains a critical open question how interhemispheric interactions in higher-order regions such as the prefrontal cortex (PFC) support working memory across the full visual space (5–7).

While V1 neural receptive fields are almost exclusively contralateral, prefrontal neurons can be tuned to either hemifield (8–10), though more neurons exhibit a contralateral preference (11,12). To explain how this organization can lead to integrated working memory across space, recent studies propose two alternative theories. In a specialized architecture, each hemisphere is predominantly responsible for contralateral information (Fig. 1a) (7). In contrast, in a redundant architecture both hemispheres represent the entire spatial field and equally guide behavior across both hemifields (Fig. 1b) (5). Each architecture helps explain conflicting empirical findings and also offers intuitive functional advantages.

**Figure 1:**
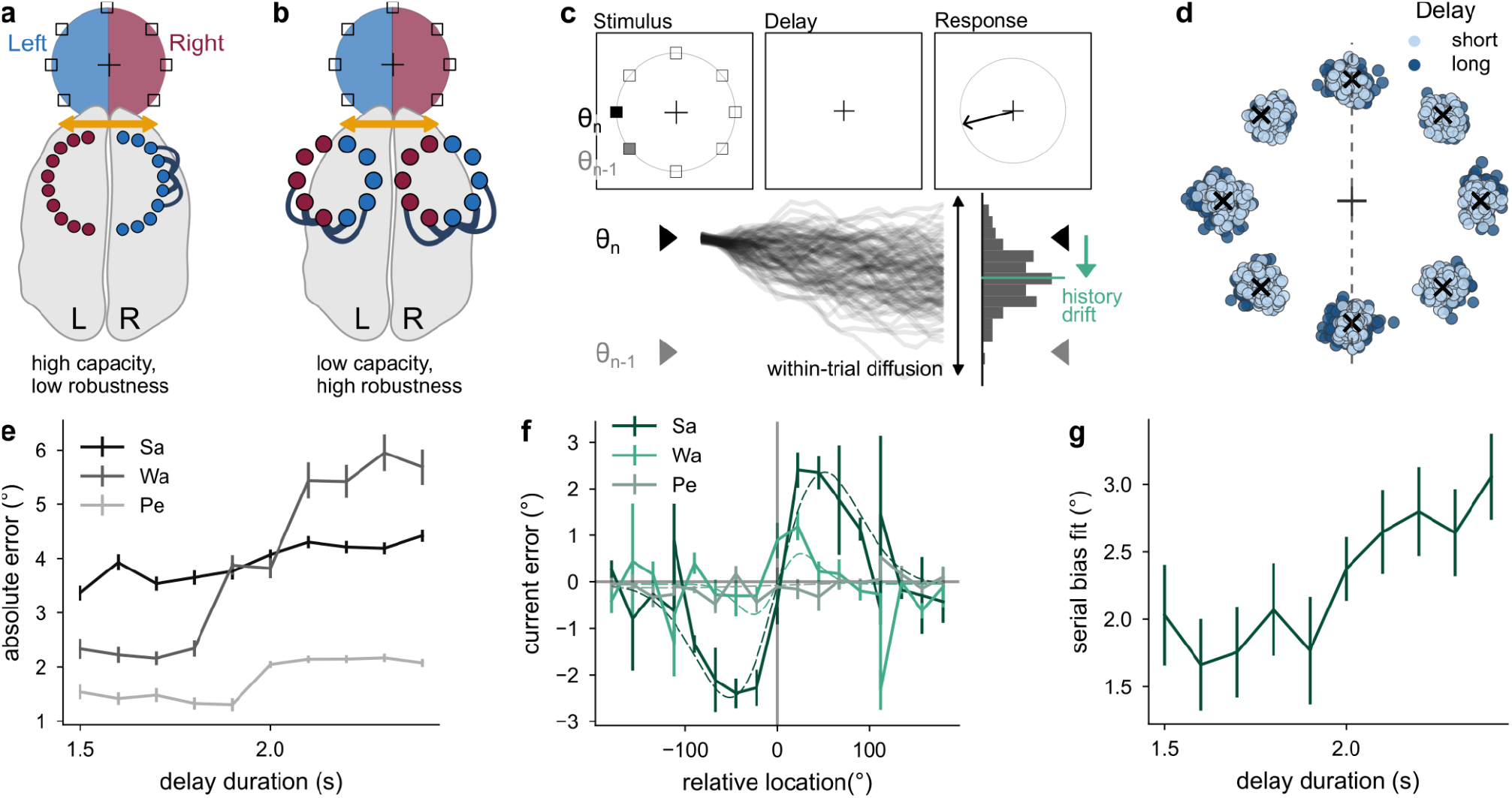
Competing interhemispheric models and task behavior. a) Specialized model: Each hemisphere primarily represents contralateral memoranda, supporting high capacity, but low robustness. Blue/red circles: left/right-preferring neurons; arrow: interhemispheric communication. b) Redundant model: Both hemispheres represent both hemifields, supporting robustness at the cost of capacity. c) Top: Behavioral task schematic indicating previous (θ_n-1_), current (θ_n_) stimulus locations. Ring, unfilled squares were invisible to the animal. Bottom: Errors may arise through random within-trial diffusion and across-trial history drift (towards θ_n-1_). d) Example session (monkey Sa): Responses (dots) clustered around targets (crosses). e) Absolute errors grew with delay length. Fit with ordinary least-squares (OLS) regression (absolute error ∼ delay; two-sided). Monkey Sa: β = 0.77 °/s, 95% CI [0.53, 1.0], t(7902) = 6.31, p = 2.95×10-10, N = 7904 trials across 16 sessions (non-repeated measures). Monkey Pe: β = 0.85 °/s, 95% CI [0.64, 1.06], t(7632) = 8.07, p = 7.94×10-16, N = 7634 trials across 16 sessions. Wa: β = 4.76 °/s, 95% CI [4.21, 5.31], t(2644) = 16.95, p = 3.08×10-61, N=2646 trials across 9 sessions. Across monkeys: β = 0.63 °/s, 95% CI [0.46, 0.79], t(18182) = 7.56, p = 4.38×10-14, with N = 18184 trials across 41 independent sessions of n = 3 monkeys. f) Error in relation to relative distance of previous and current target (serial dependence) showed idiosyncratic differences between animals: strong for monkey Sa, weak for Wa, absent for Pe. For each monkey we fit an OLS model (error ∼ DoG1; two-sided, uncorrected) with fits shown as dashed lines. Monkey Sa (σ = 0.9 rad): β = 2.42º, 95% CI [2.23, 2.61], t(7902) = 24.83, p = 4.05×10-131, N = 7904 trials across 16 sessions. Monkey Wa (σ = 0.45 rad): β = 0.66º, 95% CI [−0.02, 1.33], t(2644) = 1.92, p = 0.055, N = 2646 across 9 sessions. Monkey Pe (σ = 2.15 rad): β = 0.06º, 95% CI [−0.02, 0.14], t(7632) = 1.4, p= 0.16, N = 7634 trials across 16 sessions. g) Serial dependence fit grew with delay for monkey Sa, but not for the other monkeys (OLS model, error ∼ DoG1 × delay; two-sided). Interaction term (DoG1:delay) for Monkey Sa: β = 1.03 °/s, 95% CI [0.41, 1.65], t(7900) = 3.26, p = 0.0011, N = 7904 trials across 16 sessions. Monkey Pe: β = 0.22 °/s, 95% CI [−0.12, 0.55], t(7360) = 1.28, p = 0.2, N = 7634 trials across 16 sessions. Monkey Wa: β = 0.06 °/s, 95% CI [−3.17, 3.29], t(2642) = 0.03, p = 0.97, N = 2646 trials across 9 sessions. Error bars represent ±1 s.e.m.

A useful analogy is that of data storage in two hard drives. The strategy of maintaining unique files in each drive (specialization) maximizes storage capacity, while duplicating files across hard drives (redundancy) ensures robustness against failure. For everyday data, capacity may take precedence, but for critical files, backup is essential. Similarly, the optimal neural architecture for spatial working memory may depend on behavioral demands.

Evidence for the specialized architecture arises from behavioral and neural findings in multi-item working memory studies in human and non-human primates. Behaviorally, this is reflected in the bilateral field advantage: the ability to memorize more items (higher capacity) when items are located in different visual hemifields compared to when items are confined to one hemifield (13–20). For example, humans can attentionally track up to eight items presented bilaterally, but only four when presented within a single hemifield (13), suggesting independent storage slots for each hemisphere.

This hemispheric specialization is further supported by electrophysiological evidence, where the prefrontal neural code for a memorized item is only diminished when additional items are presented in the same, but not the opposite hemifield (15,19). Similarly, distractors affect memorized items more strongly when they are presented in the same rather than the opposite hemifield, in both behavior and electroencephalography recordings (21). Additionally, working memory representations transfer between prefrontal hemispheres following gaze shifts across the vertical meridian (22). Taken together, these studies suggest a model of hemispheric specialization with largely independent working memory resources in each hemisphere, which selectively communicate (7).

Conversely, optogenetic studies in mice suggest a redundant architecture, since short-term memory can recover from unilateral but not bilateral prefrontal perturbations (5,23,24). Additionally, frontal neural receptive fields encompass the full visual space, albeit with a contralateral preference, in mice (23,25,26) and monkeys (8–11,27). These findings suggest that each hemisphere is sufficient to store memories and guide behavior across the full visual space (Fig. 1b), challenging the specialized perspective. It is unclear if this conflicting evidence corresponds to differences in task design, brain regions, or animal species, or if it can be integrated into a unified framework.

A fundamental conceptual tool to interpret neural recordings in spatial working memory are bump attractor networks (28–30). These models can replicate working memory features such as delay-dependent precision (29,31), history effects (32–35), location-dependent distraction (21,29), feature-binding through synchronization (36), and limited multi-item working memory capacity and interference based on within-network attractor competition (37–41). This model thus constitutes a prime tool to address the computational basis of spatial working memory, which we leverage here to clarify the architecture underlying interhemispheric spatial working memory.

In this study, we mainly focus on two of these key experimental observations that characterize working memory. First, we analyze within-trial memory diffusion, known to explain response inaccuracies of working memory precision through random diffusion in persistent neural activity (29,31). Second, we study across-trial history drift, explaining the attraction of current memory reports to the previous one (serial dependence) (42–49) based on synaptic plasticity traces (32–35,50). We selected these effects as they operate on different timescales and involve distinct mechanisms, providing complementary insights into the dynamics of interhemispheric memory processing.

We investigated the role of prefrontal hemispheres in these phenomena by analyzing simultaneous bilateral recordings from monkey PFC during a visuo-spatial working memory task. Similar to previous work (8–11), our neural analyses showed that working memory across visual space was stably and simultaneously represented by both prefrontal hemispheres in a lateralized manner. Importantly, activity from either hemisphere could guide bilateral behavior, consistent with redundant memories. Neural decoding errors were weakly, but significantly correlated across hemispheres, suggesting weak interhemispheric connections that served to coordinate the working memory representations. We modeled this architecture through two redundant bump attractor networks of spiking neurons, building on recent multi-network frameworks (21,36,51–53), which was consistent with findings of memory diffusion and history drift. Importantly, simulations revealed that this architecture balances the complementary advantages of storage redundancy and specialization across different task complexities, which we validated in behavioral data from a multi-item working memory human study. Taken together, our results offer a unified network architecture that reconciles previously conflicting findings.

## Results

### Quantifying task-related behavior: Working memory precision and serial dependence

We investigated the prefrontal hemispheric organization of spatial working memory by analyzing behavior and bilateral population recordings from the dorsolateral prefrontal area 8Ar, a region important for spatial working memory, of three macaque monkeys performing a visuo-spatial oculomotor delayed response task (54,55). While the monkeys were fixating, a stimulus appeared in one of 4, 8, or 16 possible locations (depending on the session), which had to be remembered for a variable delay period (1.5-3 s) and reported through a saccade to the remembered location (Fig. 1c, top).

We analyzed two main behavioral effects: working memory precision, defined as the variability in memory recall of stimulus locations, and serial dependence, the systematic attraction of current reports towards previously remembered items. These behavioral effects have been mechanistically linked to the diffusion of persistent activity (29,31) and drift based on synaptic plasticity (32,33) through model-driven analyses of electrophysiological recordings (Fig. 1c, bottom) and can offer complementary insights on working memory processing at different time scales.

Monkeys exhibited good accuracy overall, but absolute precision errors significantly increased with longer delays (Fig. 1d,e), as previously reported (56–58), suggesting memory diffusion (31).

We also observed serial dependence (SD, Fig. 1f), replicating findings in both humans (32,34,42–50,59–66) and monkeys (32,56,67). We measured the effect size by fitting the response error using a first-order derivative of Gaussian to fit the serial dependence curve (DoG1 curve; hyperparameters in Supplementary Fig. 7a-c; see Methods). We found varying degrees of serial dependence across monkeys (Fig. 1f), akin to known idiosyncratic differences in humans (50,68,69). The monkey with the strongest serial dependence showed a clear build-up of serial dependence with longer delays (Fig. 1g), as previously reported (34,42,45,67). Both precision and serial dependence were comparable for stimuli presented in the left and right hemifield (Supplementary Fig. 1).

### Each hemisphere stably represented the full visual space with a contralateral preference

While the monkeys performed this task, neural activity was recorded from bilateral 96-channel Utah arrays in area 8Ar of the PFC (see Methods). We recorded an average of 139.2 ± 3.8 multi-units in each of the 41 sessions across both hemispheres, and excluded one session of monkey Pe from the neural analyses, due to a shorter stimulus duration. We found tuned persistent activity in trial-averaged data collected during the delay period of the task (example multi-unit in Fig. 2a), as previously reported (11,31,32,70). We leveraged the large population recordings to obtain a single-trial view of the population activity by showing the evolving firing rates of multi-units sorted by their memory fields (cross-validated, see Methods). In an example session with 158 multi-units, we observed clear, tuned persistent activity in some individual trials from stimulus onset until the response (example trial in Fig. 2b, different stimulus locations Supplementary Fig. 2, all trials from this session Supplementary Fig. 3). As expected, we saw clear tuning during the delay in the trial-averaged activity (example session in Fig. 2c).

**Figure 2:**
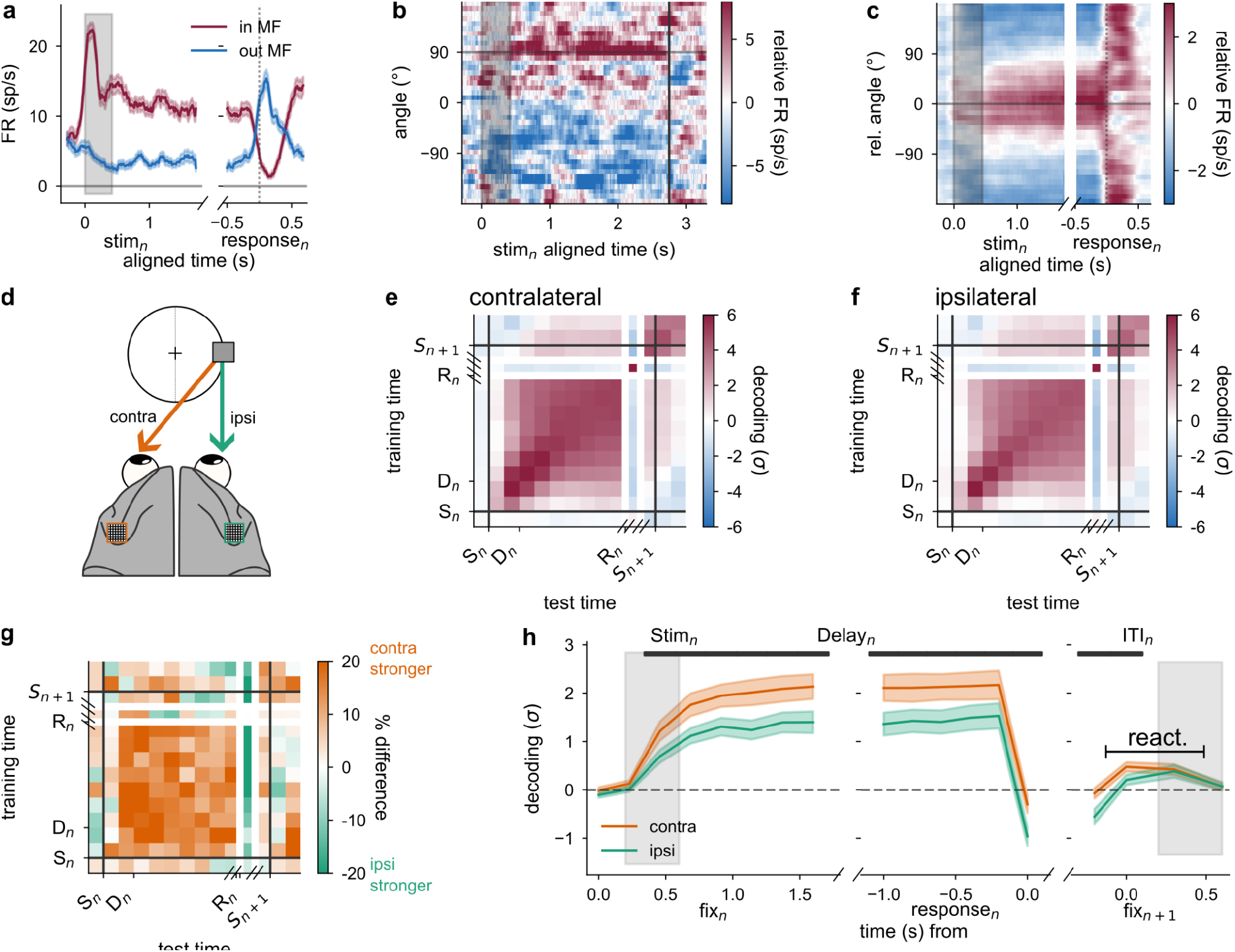
Hemispheres represent the full visual field with a contralateral preference. a) Example multi-unit: Firing rate averages of cues in/out of the memory field. 59.13% (3375) of 5708 delay-selective multi-units (Sa: 88.4% (1950 of 2206 selective units), Pe: 28.1% (666 of 2370 selective units), Wa: 72.23% (705 of 976 selective units), bootstrapped test, see Methods, differences likely due to sampling differences of the implanted arrays). b) Firing rate relative to each neuron’s average (see Methods) sorted by preferred location (estimated on left-out trials, Methods) from stimulus start (gray; stimulus at 90°) to response (dotted line). c) Session average of panel b, after recentering around the stimulus location. d) Schematic of the bilateral arrays in area 8Ar in a right stimulus trial (square), rendering the left (right) hemisphere contralateral (ipsilateral). e) Cross-temporal decoder accuracy in contralateral trials, expressed as z-scored distance from shuffle baseline (σ, Methods). Task events: stimulus (S_n_), delay (D_n_), response (R_n_), next stimulus (S_n+1_). White lines indicate discontinuity because of variable delays; black lines indicate stimulus presentation. f) same as panel e for ipsilateral trials. g) Percent lateralization in decoding ((contra-ipsi)/contra×100), showing stronger contralateral coding (14.94% stronger in the delay). h) Delay decoder (trained on average delay activity) showed lateralization during the delay. Paired two-sided t-test between contralateral and ipsilateral delay decoding (7 response-aligned independent 200ms bins): t(33) = 10.66, p = 3.20×10^-12^, mean = 0.7, 95% CI [0.57,0.84]. At the subsequent trial’s fixation (fix_n+1_) both hemispheres showed reactivations similarly. One-sample two-sided t-test against 0 of the average reactivation decoding (3 independent 200ms bins) revealed ipsilateral reactivations: t(33) = 2.54, p = 0.016, mean = 0.22, 95% CI [0.04, 0.4] and contralateral reactivations: t(33) = 4.86, p = 2.76×10^-5^, mean = 0.32, 95% CI [0.19, 0.46]. Paired two-sided t-test between contralateral and ipsilateral reactivations: t(33) = 1.8, p = 0.08, mean = 0.1, 95% CI [−0.01, 0.21]. All tests were computed across N = 34 independent sessions from n = 3 monkeys (6 sessions excluded due to insufficient ipsilateral/ contralateral targets for shuffle control, 1 session excluded due to shorter stimulus duration). Error bars represent ±1 s.e.m.

We investigated the extent of hemispheric lateralization across sessions by linearly decoding the stimulus location from the population activity in each hemisphere (see Methods; firing rates in Supplementary Fig. 4). Decoders were trained on 80% of the trials at each time step and the mean-squared error between predictions and stimulus was computed on contralateral and ipsilateral left-out trials separately (schematic Fig. 2d). We compared the mean-squared error with a stimulus-shuffled decoder (decoding: z-score to shuffle; see Methods) and found a stable delay code in both contralateral (Fig. 2e) and ipsilateral (Fig. 2f) trials. Contralateral outperformed ipsilateral decoding by around 15% (averaged delay difference, Fig. 2g).

In the following, we used a decoder trained on the average neural activity during the delay period (Fig. 2h, Delay Decoder, Methods) to focus on memory encoding rather than other task parameters, such as motor responses (71). This Delay Decoder revealed a reactivation of memory-related activity during fixation of the upcoming trial (line at fix_n+1_, Fig. 2h), previously shown to correlate with serial dependence strength (32,50,72–74). We could not replicate this effect here, which may be due to varying, shorter inter-trial intervals, variable delays or other differences in the experimental design. Notably, reactivations were present in both hemispheres (Fig. 2h), indicating little lateralization in their occurrence. Based on previous proposals in a single hemisphere (32,50), this suggests that each hemisphere contains enough persistent activity to produce reliable synaptic traces which can be reactivated at the start of the next trial.

### Both hemispheres equally reflected trial-by-trial behavioral variability with weakly correlated memory representations

To address how PFC activity related to behavioral responses, we correlated the signed (clockwise, counterclockwise) neural decoder errors with the corresponding signed behavioral response errors (31,56). A positive correlation during the delay would indicate that the multi-units can predict behavioral imprecisions before they occur. Furthermore, a steady increase throughout the delay suggests slowly diffusing memories as proposed by attractor dynamics (31,56). Indeed, we found steadily increasing correlations when pooling multi-units from both hemispheres (Fig. 3a).

To understand how hemispheric memory representations are combined to provide unified behavior, we separately trained decoders on each hemisphere and performed the same analysis. We found that both the left and right hemispheres showed significant neural-behavioral correlations (Fig. 3b, Supplementary Fig. 5b). Similarly, both ipsilateral and contralateral trials significantly correlated with behavior (Fig. 3c, Supplementary Fig. 5a). Interestingly, when we exclusively considered trials in which hemispheric predictions were in opposite directions (clockwise vs. counterclockwise), neither hemisphere dominated the correlation with behavior (Supplementary Fig. 5c). Therefore, even though the memory code was more precise in contralateral trials (Fig. 2g), both hemispheres seemed of similar importance for the behavior. Thus, in combination with the strong ipsilateral decoding (Fig. 2h), our analyses did not support the specialized model (Fig. 1a), but rather pointed towards a redundant architecture (Fig. 1b), with both hemispheres encoding stimuli and guiding behavior in the full visual field.

**Figure 3:**
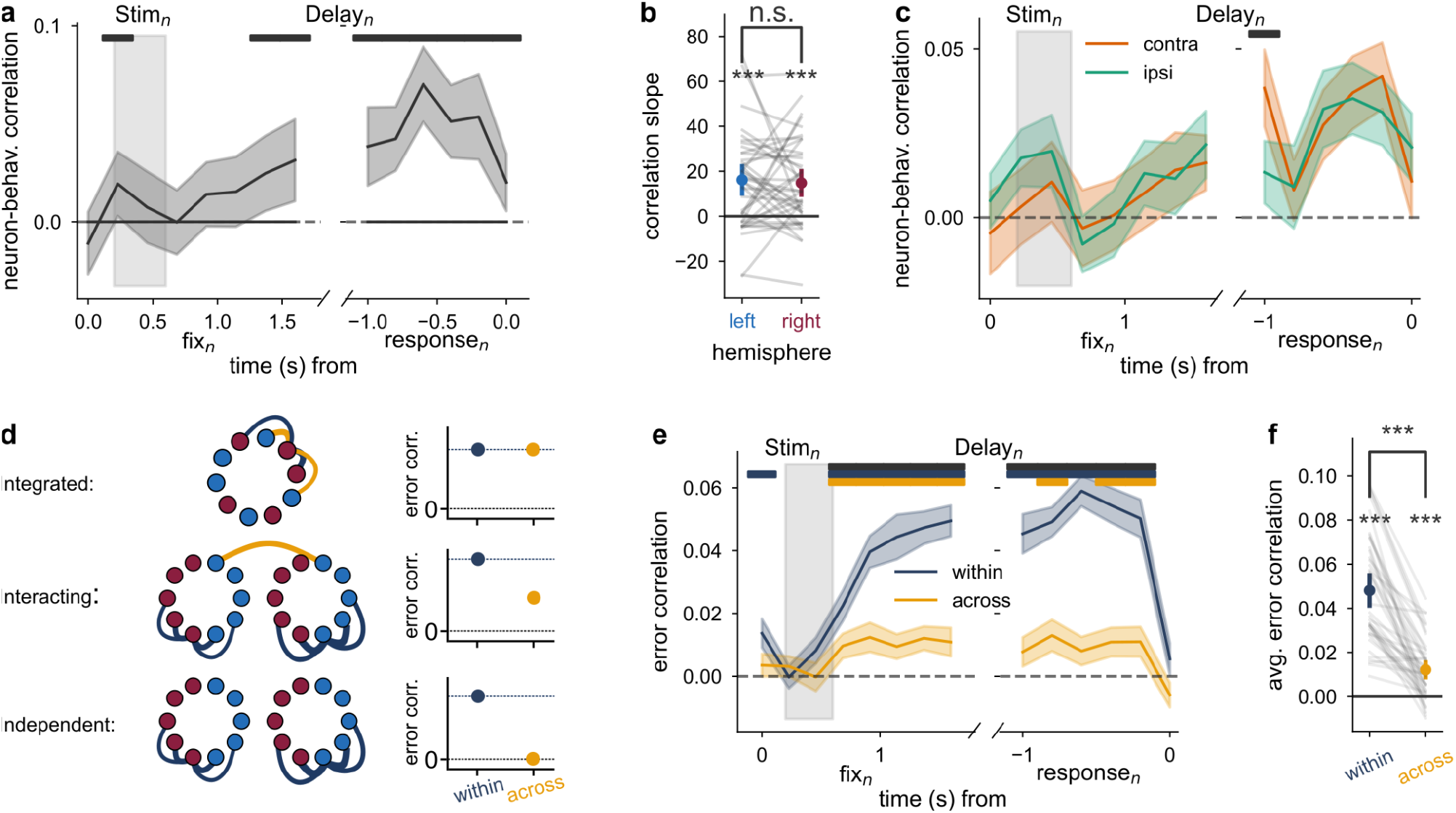
Memories were stored redundantly in both hemispheres with interacting representations. a) Correlation of decoder and behavioral response errors. Slopes were fit on 7 response-aligned independent 200ms bins through linear regression for each of the independent N = 40 sessions across n = 3 monkeys. One-sample two-sided t-test against 0: t(39) = 2.4, p = 0.021, mean = 0.02 1/s, 95% CI [0.0, 0.05]. Error bars are 95% C.I. b) Same as panel a, but showing no significant difference in slope for left and right hemisphere predictions. Paired two-sided t-test: t(39) = 0.38, p = 0.71, mean = 0.0 1/s, 95% CI [−0.01, 0.01]. c) Same as panel a, but for contralateral vs ipsilateral trials. Paired two-sided t-test: t(39) = 0.39, p = 0.7, mean = 0.0 1/s, 95% CI [−0.01, 0.01]. d) Schematic of different network architectures (left) predicting different across-hemisphere error correlations (right). e) Decoder prediction error correlations within and across hemispheres across time. f) Delay-averaged (average of 7 response-aligned independent bins) error correlations from panel e, individual sessions are grey lines. Error correlations within and across hemispheres differed significantly from zero. One-sample two-sided t-test to 0, within: t(39) = 12.25, p = 6.08×10^-15^, mean = 0.05, 95% CI [0.04,0.06]; across: t(39) = 5.6, p = 1.87×10^-6^, mean = 0.01, 95% CI [0.01,0.02]. Correlations within each hemisphere were stronger than across hemispheres. Paired two-sided t-test t(39) = 9.66, p = 6.80×10^-12^, mean = 0.04, 95% CI [0.03,0.04]. All t-tests are performed across N = 40 independent sessions from n = 3 monkeys. Error bars are ±1 s.e.m. unless otherwise specified. Top bars denote significant differences of one-sample two-sided t-test against 0 (colors, black in panel a) or paired two-sided t-tests between samples (black).

We further aimed to specify the interhemispheric organization underlying the redundant working memory network. A redundant architecture can range from being so tightly coupled that the hemispheres are essentially integrated (Fig. 3d, top) to two uncoupled hemispheres that each represent the full visual space (Fig. 3d, bottom). These alternatives have different predictions for memory diffusion in each hemisphere. If both hemispheres are integrated, memories would be perfectly mirrored with identical encoding errors (Fig. 3d, top). On the other hand, unconnected hemispheres would diffuse independently leading to uncorrelated memory errors (Fig. 3d, bottom, right).

We used the decoder prediction errors as a proxy for the memory imprecisions of each hemisphere. Specifically, we repeatedly trained four decoders on random halves of the multi-units recorded from each hemisphere (sub-hemispheres: left_A_, left_B_, right_A_, right_B_) and obtained their prediction errors. We correlated the prediction errors either within (within: left_A_ vs. left_B,_ and right_A_ vs. right_B_) or across hemispheres (across: left_A_ vs. right_A_, and left_B_ vs. right_B_). We found that errors were more strongly correlated within than across hemispheres (Fig. 3e,f), suggesting separate functional networks for each hemisphere, which separately accumulate memory errors.

However, interhemispheric decoder error correlations were significantly above zero (Fig. 3e,f, across), suggesting either connections across hemispheres or a retinotopically-tuned common input to both hemispheres. To address a plausible source of common input, we considered uninstructed eye movements while monkeys fixated (e.g., microsaccades). Such movements could induce memory imprecisions and also reflect brain-wide arousal signals (54,75,76), accounting for two types of common input. Indeed, eye gaze locations correlated with the memorized stimulus (Supplementary Fig. 6a-b) (75,77–79). However, gaze errors did not correlate with neural prediction errors (Supplementary Fig. 6b), making it unlikely that the eye movements provided common input related to memory errors. Indeed, even when removing uninstructed eye movements from the decoder analysis (see Methods), the same pattern of across-hemisphere error correlations remained (t-test across: t = 4.2, p = 0.00015, Supplementary Fig. 5c, d). Our analyses therefore suggest that the observed interhemispheric error correlations arise from interacting hemispheres (Fig. 3d, middle) and not from a gaze confound.

### Tuned across-area connections led to shared memory diffusion in a working memory model

To understand how redundant, interacting hemispheres can support the observed interhemispheric working memory effects, we implemented the proposed architecture (Fig. 3d, middle) in a computational network model (Fig. 4a). We used the bump-attractor framework (29), which can model memory diffusion consistent with the observed neural-behavior correlations (Fig. 3a) (31).

**Figure 4:**
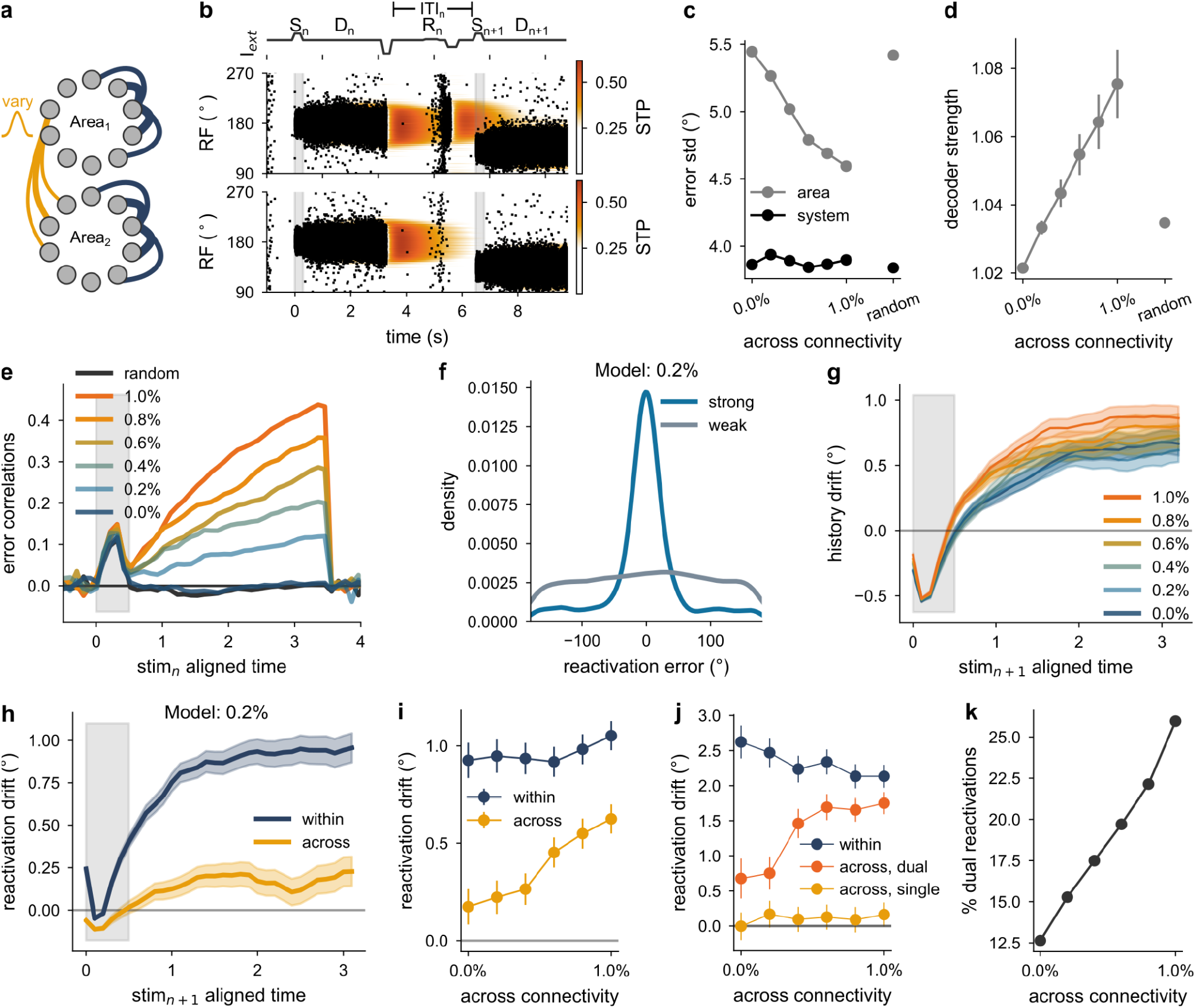
Tuned across-area connectivity increased coordination of within-trial memory diffusion and across-trial memory drift. a) Model schematic: Two bump-attractor networks with short-term plasticity (STP) and varying across-area connectivity (yellow). b) Example simulation of two consecutive trials. Only Area_1_ but not Area_2_ showed a reactivation. Top: External stimulus strength. S, D, R and ITI mark stimulus onset, delay, reactivation, and inter-trial interval, respectively. Middle and Bottom: Spiking activity (dots) and STP dynamics (heatmap) for Area_1_, Area_2_, respectively. c) Increasing tuned inter-area connectivity reduced response errors of each area, while system-wide (average of both areas) response errors remained consistently low. d) Tuned connectivity enhanced memory stability (decoding strength). e) Only models with tuned across-area connections exhibited inter-area error correlations. f) Strong reactivation trials (38% of trials) showed a tighter replication of the previous stimulus location. g) Neural delay activity increasingly drifted towards the previous stimulus (history drift; estimated using Dog1 fit, Methods) over time. h) Drift of the neural delay activity towards reactivations within the same hemisphere was stronger than across hemispheres (exemplary for the 0.2% model). i-j) Stronger across-area connectivity increased shared reactivation drift (taken at the end of the delay), particularly in trials with dual reactivations. k) The percentage of dual reactivations increased with connectivity strength. Total of N = 60,000 simulations. Error bars represent ±1 s.e.m.

Specifically, we modeled each hemisphere as a bump-attractor network of spiking excitatory and inhibitory leaky-integrate-and-fire neurons covering the full visual space. Brief external stimuli were delivered to each ring (henceforth called area) at the beginning of each trial (Fig. 4b, S_n_, S_n+1_), which were maintained in each area network through recurrent excitation after the stimulus offset, leading to tuned persistent activity (Fig. 4b, D_n_, D_n+1_).

We studied the effects of different interhemispheric interactions by manipulating the strength and structure of excitatory connections between the two areas (Fig. 4a), expressed as a percentage of the excitatory coupling strength within each area (see Methods). In particular, we compared unconnected areas (0.0%), untuned connected areas (random, 1.0% of within connectivity), and varying levels of tuned across-area connections, where neurons with similar receptive fields were more strongly connected across areas (0.2% - 1.0%) in a total of N = 60,000 simulations.

Irrespective of connectivity type, memory precision was higher when responses were based on the averaged population activity of both areas (system), compared to on a single area (area, Fig. 4c), highlighting that hemispheric redundancy improved memory performance. Precision within each area improved with tuned across-area connectivity (Fig. 4c, grey) due to enhanced population tuning and therefore stability of each area’s memory bump (Fig. 4d). The system-wide memory precision on the other hand remained stable across connectivity types (Fig. 4c, black),

To relate the data to the model, we asked which model can replicate the correlated memory errors observed in the neural analyses (Fig. 3e). Neither the uncoupled model (Fig. 4e, 0.0%) nor the model with random 1.0% connectivity (Fig. 4e, random) produced across-area error correlations. Only increasing degrees of tuned connectivity increased error correlations (Fig. 4e), suggesting that the prefrontal hemispheres are connected via tuned interhemispheric connections, in line with noise correlation analyses in these data (Fig. S2 in (54)).

Together, our simulations show that redundancy enhances memory precision and suggest that interhemispheric connections in the monkey PFC are tuned, supporting local memory stability.

### Increasing across-area connectivity in a working memory model increased shared history drift

We further used this model to investigate memory dynamics at longer timescales, namely across-trial memory drift underlying serial dependence. We could analyze this in the model, since within-area excitatory connections were facilitating (80), increasing synaptic efficacy after pre-synaptic spiking (short-term plasticity, see Methods) and leading to serial dependence in this task (32–35,50).

Guided by previous work with a single area (32–34,50), we simulated sequential trials separated by an activity-silent inter-trial interval (Fig. 4b, ITI_n_). Towards the end of the inter-trial interval, a nonspecific excitatory current (a reactivation current (R_n_), Fig. 4b) was applied to both networks, generating a bump reactivation in 38% of trials. Due to synaptic plasticity, reactivations were typically centered at the location of the previous stimulus (Fig. 4f) and further replenished the synaptic plasticity (heatmap between R_n_ and S_n+1_, Fig. 4b, top), leading to stronger attraction to the previous memory in the upcoming trial (32). Furthermore, we modeled adaptation, a repulsive sensory history effect originating in lower-level visual areas (49,81–83), by shifting stimulus inputs away from the previous trial’s location (34,35) (see Methods). Overall, we found a delay-dependent increase in serial dependence from an initial repulsion to a later attraction as observed in human experimental data (34,42,45) for all connectivity strengths (Fig. 4g).

To clarify the impact of across-area connectivity strengths on history drift, we analyzed how the delay-period mnemonic activity (Fig. 4b, D_n+1_) in each area drifted towards the previous fixation-period reactivations (Fig. 4b, R) of the same area (within) or across areas. In the weakest-tuned model, the attraction was much stronger within than across areas (Fig. 4h), but increasing the inter-area connectivity increased shared history drift (Fig. 4i). This effect was driven by trials in which reactivations occurred concurrently in both hemispheres (Fig. 4j, dual), with the history drift remaining private when reactivations only occurred in a single area (Fig. 4j, single). Therefore, shared history drift was driven by an increased amount of dual reactivation trials as inter-area connectivity increased (Fig. 4k).

In sum, our simulations showed that reactivations and with them across-trial history drift were affected by increasing across-area connectivity strength. Notably, history effects could be transferred across areas only if across-area connectivity strength was high and simultaneous reactivations occurred in both areas (dual trials). Based on these insights from the model we next evaluated inter-hemispheric history effects in the neural data.

### Each prefrontal hemisphere had private history effects, suggesting weak interhemispheric connectivity

We first confirmed that neural activity during memory maintenance in monkey PFC was biased towards previous stimulus locations. This across-trial history drift is a neural correlate of serial dependence previously found using non-invasive recordings in humans (43,49,66,83) and is parsimoniously explained by bump-attractor models (Fig. 4g) (32).

Neural decoder errors were biased towards previous stimuli (Fig. 5a), mirroring behavioral serial dependence (Fig. 1f). Interestingly, the neural code was repelled from the previous stimulus during stimulus presentation but became attracted at the end of the delay period (Fig. 5a, example monkey Sa). To obtain a time-resolved estimation of the neural serial dependence, we fit a first-order derivative-of-Gaussian curve to the predicted errors at each time (dashed lines in Fig. 5a; hyperparameters in Supplementary Fig. 7d-f; see Methods) and observed a gradual change from repulsion to attraction throughout the delay (Fig. 5b). Notably, the neural history drift scaled with the behavioral serial dependence strength across monkeys (Supplementary Fig. 8a), though more subjects are needed to establish a correlation at the population level. From now on, the analyses focus on the monkeys with positive behavioral and neural serial dependence (n=2, Monkeys Sa, Wa).

**Figure 5:**
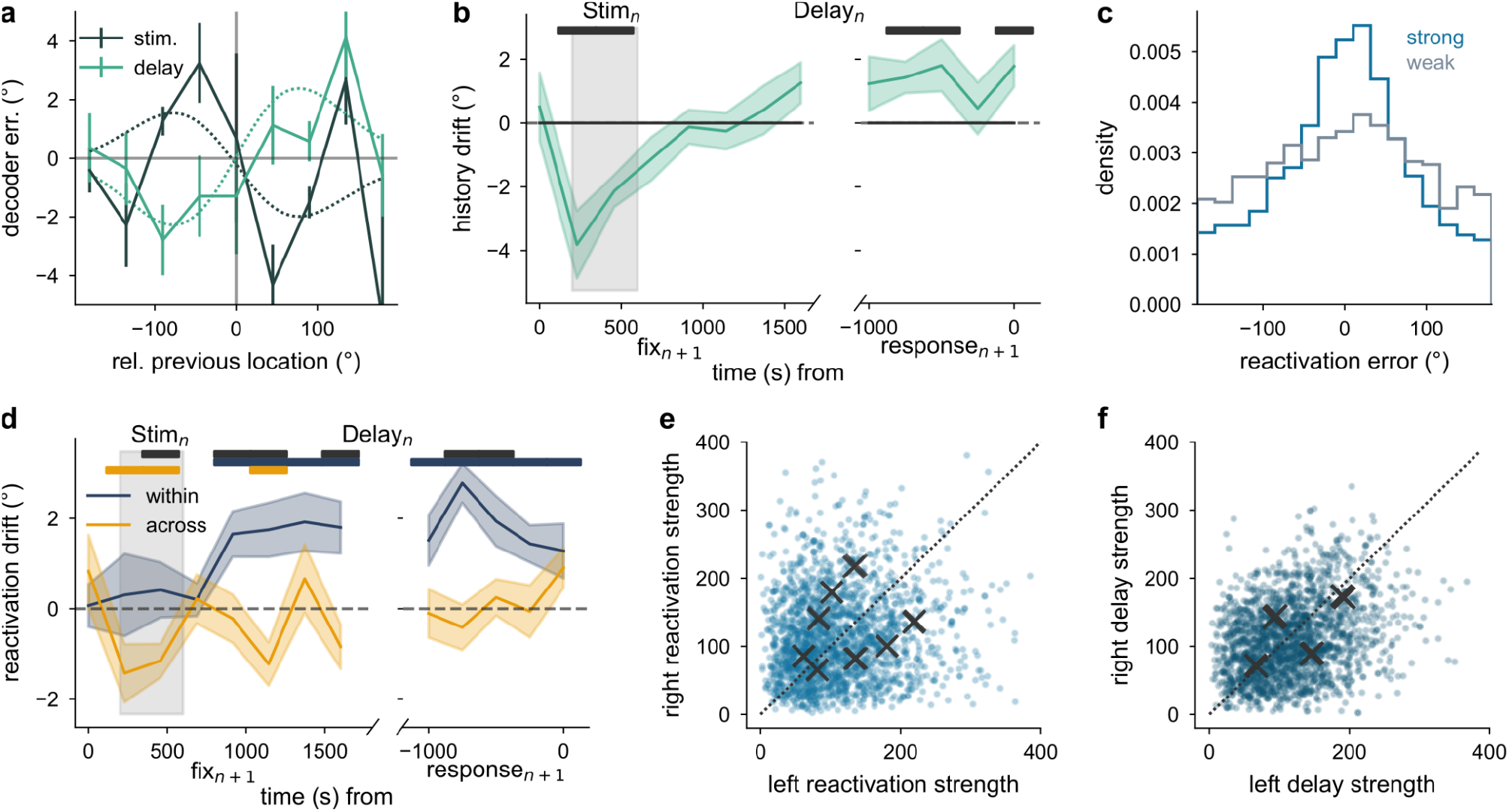
Private across-trial history drift in monkey PFC indicates weak but specific interhemispheric coupling. a) Example monkey Sa: Session-averaged decoder error over relative distance of previous and current stimulus locations. Dashed lines indicate OLS model fit for ‘error ∼DoG1’ (see Methods). Stimulus: *β* = −1.76°, 95% CI [−2.37, −1.16], t(7902) = −5.71, p = 1.20×10^-8^; Delay: 2.33°, 95% CI [1.4, 3.25], t(7902) = 4.93, p = 8.20×10^-7^ for N = 7904 independent trials of 16 sessions of n=1 monkey (Sa). b) Time-resolved neural history drift (fit from panel a at each time) across the n = 2 monkeys with significant behavioral serial dependence (monkeys Sa, Wa) from stimulus start to stimulus-aligned delay end (9 independent 200ms bins) for N = 25 sessions. One-sample two-sided t-test to 0: t(24) = 8.73, p = 6.46×10^-9^, mean = 3.0 °/s, 95% CI [2.6, 4.2]. c) Example monkey (Sa): Precision of reactivation location (average of fixation to stimulus) relative to the previous stimulus location for strongly/ weakly reactivated trials (cut-off at 30% of each session’s decoder strength; see Methods). d) Attraction towards the previous reactivation location was private within each hemisphere. One-sample two-sided t-test showed average delay reactivation drift (7 independent response-aligned 200ms bins) within each area t(24) = 6.86, p = 4.27×10^-7^, mean = 1.86°, 95% CI [1.3, 2.42] but not across areas t(24) = 0.21, p = 0.83, mean = 0.05°, 95% CI [−0.42, 0.51]. Paired two-sided t-test showed a significant difference between conditions t(24) = 5.55, p = 1.05×10^-5^, mean = 1.81°, 95% CI [1.14,2.49]. Test performed over N = 25 sessions from n = 2 monkeys with significant behavioral serial dependence. e) Distribution of decoded reactivation strengths in the left and right hemispheres and fit class means (crosses, four-class Gaussian Mixture Model (250 repeats)). f) Same as panel e for delay prediction strengths. Error bars represent ±1 s.e.m.; Top bars indicate significance of one-sample two-sided t-test to 0 (colors, black in panel b) or paired two-sided significance between conditions.

We tested whether this neural history drift was private to each hemisphere. For this, we computed the attraction of the neural code towards the previous reactivation location decoded from either the same or the opposite hemisphere, as in the model (Fig. 4h). First, we verified that reactivations could be extracted using a Delay Decoder (see fix_n+1_ in Fig. 2h) and were tuned to the prior stimulus location in strongly reactivated trials (Fig. 5c). Then, we assessed how the neural code drifted towards the prior reactivations (DoG1 fit, hyperparameters in Supplementary Fig. 7g-i; see Methods). We exclusively found memory drift to reactivations occurring in the same hemisphere (within), but not the opposite hemisphere (across, Fig. 5d). Across-hemisphere drift was somewhat weaker than in the simulations, probably reflecting either lower incidence of dual reactivations or large noise levels, hiding a weak effect. Based on our model simulations, this finding of hemisphere-specific history drift suggests weak interactions between the hemispheres (Fig. 4h,i).

As in the model (Fig. 4j), we also investigated how different reactivation types affected the reactivation bias. We split the data into null, single and dual reactivation trials based on a 30% reactivation strength threshold for each hemisphere (Supplementary Fig. 8b). We observed that in null trials no significant reactivation bias emerged, whereas both single and dual reactivation trials showed within- but not across-hemispheric attraction (Supplementary Fig. 8 c,d). These findings pointed towards a model with very weak interhemispheric interactions (Fig. 4j, e.g. 0.2%).

The model further predicted that private reactivation drift depends on most trials exhibiting reactivations in a single hemisphere (Fig. 4i-k). We tested this prediction using a four-class Gaussian Mixture Model (GMM; see Methods) assuming four possible reactivation patterns: null, single_left_, single_right_ and dual reactivations. Across 250 runs with different parameter initializations, the GMM produced eight distinct class means, all located away from the diagonal in the decoder-strength space (Fig. 5e). This suggests two key points: (1) the underlying distribution is not well explained by four classes, as different initial conditions caused class means to shift; and (2) dual reactivations were rare, as they would appear on the diagonal in this graph.

We further found that a three-class GMM consistently identified three off-diagonal class means across all 250 re-runs, accounting for null, single_left_ and single_right_ reactivation types (Supplementary Fig. 8 e-g). Taken together, these findings suggest that three (and not four) classes underlay the distribution, with no evidence for a dual reactivation class. In contrast, during active memory-maintenance in the delay period, the four-class GMM analysis reliably estimated all four classes, including a dual class (Fig. 5f).

We conclude that the prefrontal hemispheres interacted through tuned but weak interhemispheric coupling, leading to correlated memory diffusion but private history drift and reactivations. This is particularly intriguing since it allowed the networks to interact when both hemispheres stored similar items (delay code strength correlations, Fig. 5f) but perform separately when inputs only triggered events in one hemisphere (reactivations remaining private, Fig. 5d,e).

### Reconciling competing models: Weakly connected hemispheres enabled both redundancy and hemispheric specialization

Our findings suggest weakly connected, redundant hemispheres consistent with shared memory diffusion and private history drift. These results seemingly contradict prior evidence supporting specialized hemispheres. Specifically, previous behavioral and electrophysiological studies have shown reduced memory interference when multiple items are presented across rather than within a visual hemifield, a phenomenon known as the bilateral field advantage (13–20).

However, even in our proposed redundant architecture, we observed that reactivations remained private to each hemisphere, indicating that the prefrontal hemispheres were capable of maintaining separate memory representations if they were independently activated. We hypothesized that a lateralization of inputs to the PFC from upstream areas (23,84) could preferentially activate each hemisphere for contralateral stimuli and thus promote hemispheric specialization during multi-item tasks (20,85).

To test this hypothesis, we simulated a simple multi-item working memory task in the previously proposed weakly-connected model (Fig. 4, 0.2%). We simultaneously presented two stimuli to the network, which were recalled at the end of the delay. The only modification to the previous architecture was the addition of the input lateralization. Each area was assigned a hemisphere (Area_1_ = left, Area_2_ = right) and received a weaker input for ipsilateral than contralateral stimuli (70% of contralateral input strength).

We tested two equidistant stimulus conditions. Stimuli were presented either bilaterally (one per hemifield; Fig. 6a) or unilaterally (both in the same hemifield; Fig. 6b). In bilateral trials, each area received one stronger contralateral and one weaker ipsilateral input (Fig. 6a, solid vs dashed arrows). Due to the single-item capacity of each area, this resulted in the memory maintenance of only the stronger, contralateral item in each area and therefore the successful maintenance of both items across the entire network in a specialized manner (Fig. 6c, example trial). In contrast, in unilateral trials, both stimuli were contralateral (or ipsilateral) to the same hemisphere, such that each hemisphere received two equally strong inputs (Fig. 6b). This created an equal competition between both items, causing the forgetting of a random item in each area. Crucially, both areas could forget the same item (example trial, Fig. 6d) and therefore store one item redundantly, which reduced the system-wide working memory capacity in unilateral compared to bilateral trials, replicating the well-known bilateral field advantage (Fig. 6e, 1,500 simulations).

**Figure 6:**
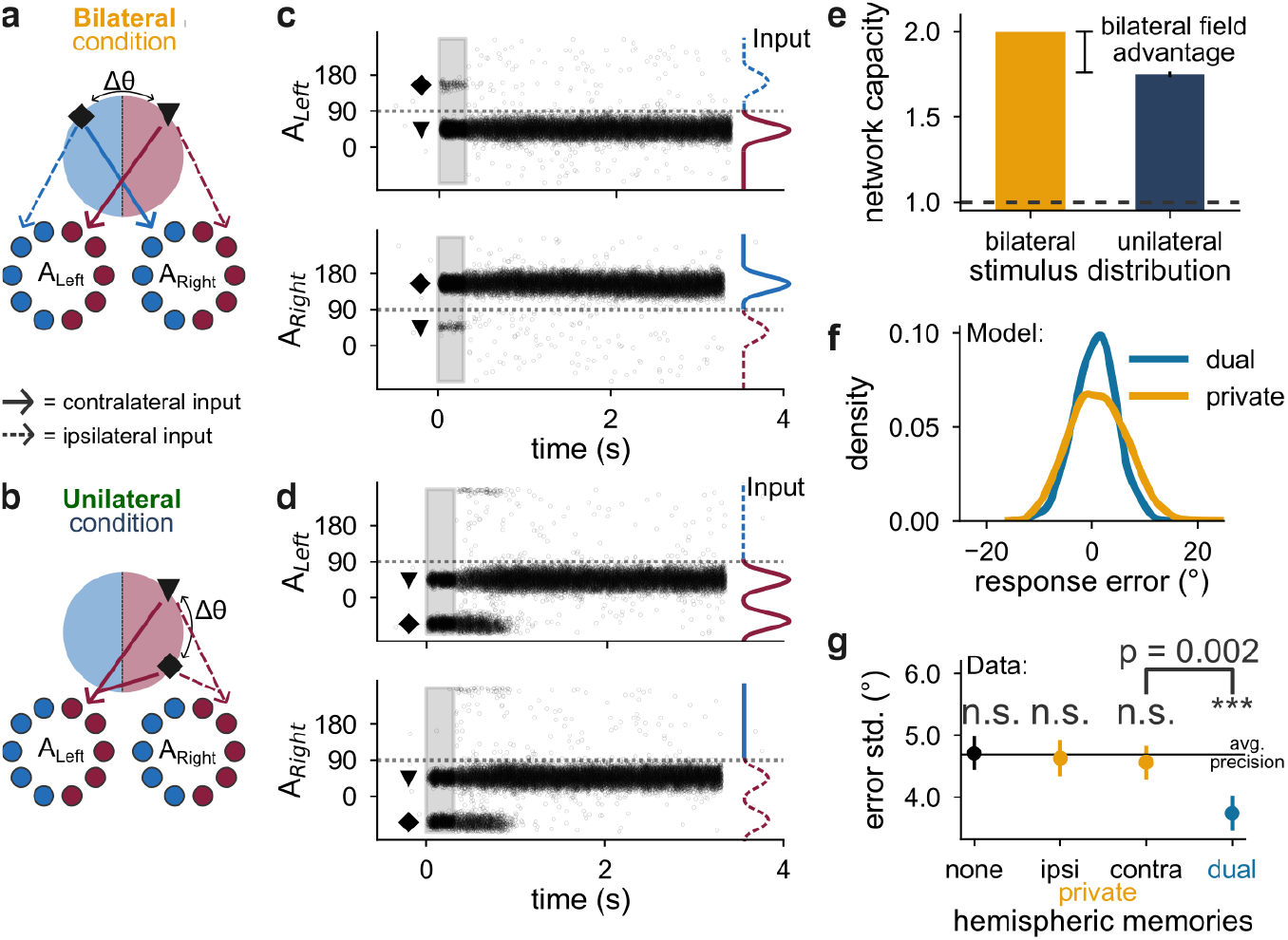
Stimulus lateralization in a redundant network model enabled hemispheric specialization in a multi-item task. a) Schematic of the model from Fig. 4 (0.2% connected) simulated (n=1500 trials) for two simultaneous bilateral stimuli (triangle, diamond), one in each hemifield with weaker ipsilateral projections (dashed arrows; 20% reduced vs. contralateral, solid arrows). b) Same schematic for the unilateral condition. Both areas received two equally strong inputs, since each hemisphere was contralateral (ipsilateral) to both items. c) Example simulation from bilateral condition. Each area retained its contralateral item, leading to both items being retained (A_Left_ retains triangle, A_Right_ retains diamond). Dashed horizontal lines indicate hemifield boundary (90°); colored Gaussians represent input strength. d) Example simulation from unilateral condition. Both areas retained the same item (diamond), causing the forgetting of the other item (triangle). e) Working memory capacity (trial-averaged number of retained items) for bilateral and unilateral stimuli (bilateral = 2 items, unilateral = 1.75 items, single=1 item capacity). f) Response precision was reduced when both items were remembered privately (bilateral condition, σ = 5.37°) compared to when one item was remembered dually (unilateral subset when one item was remembered, σ = 3.91°). g) Monkey behavioral memory errors for trials with none, private (ipsilateral, contralateral) and dual precise memory representations (20% of trials with smallest average decoder error in each session; other cut-offs in Supplementary Fig. 9). As predicted by the model, behavioral precision was improved for dual memory storage (Session-paired t-test to average precision; Dual: t(39) = −4.12, p = 1.93×10^-4^, mean = −0.95, 95% CI [−1.41,-0.48]; Private Ipsi: t(39) = −0.71, p = 0.49, mean = −0.06, 95% CI [−0.22,0.1]; Private Contra: t(39) = −1.35, p = 0.18, mean = −0.12, 95% CI [−0.31,0.06]; Dual vs. Contra: t(39) = −3.29, p = 2.12×10^-3^, mean = −0.82, 95% CI [−1.33,-0.32]); None: t(39) = 0.91, p = 0.37, mean = 0.03, 95% CI [−0.03,0.08]) for N = 40 sessions across n=3 monkeys. Error bars represent ±1 s.e.m.

Conversely, the memory precision of the network was higher when both areas redundantly stored the same item (Fig. 6f, blue) compared to when each area specialized in storing a unique item (Fig. 6f, yellow). This aligned with earlier modeling results (Fig. 4c) and led to a testable prediction for the experimental data: When both hemispheres maintain redundant memories, behavioral precision should be enhanced relative to unilateral maintenance.

To test this prediction in the data, we grouped trials according to their decoder accuracy during the late delay in each hemisphere. Accurate memories were defined as the 20% smallest absolute decoder errors in each hemisphere (alternative thresholds shown in Supplementary Fig. 9). Based on this criterion, trials were classified as none, for no accurate maintenance, private, for only one hemisphere showing accurate maintenance (yellow, ipsilateral or contralateral) and dual when both hemispheres were accurate (blue). As predicted, trials with dual prefrontal memory storage showed superior behavioral precision, reflected by a smaller standard deviation of the monkey’s response errors, compared with trials in which only one or neither hemisphere stored the item accurately (Fig. 6g).

Next, we increased the capacity of our computational network in order to develop predictions in naturalistic multi-item tasks. We modified three network parameters (narrower stimulus width, narrower recurrent weight profile, stronger excitatory-to-excitatory connections, see Methods) to increase the capacity of each area (37,40,41). Each area could store around two items, and adding a second item within a hemifield reduced firing rates more than adding an item in the opposite hemifield, consistent with previous electrophysiological findings in monkeys (Supplementary Fig. 10a) (19).

We repeatedly (n=6000 trials) stimulated the network below (load 2) and at capacity (load 4) to mimic the conditions of spatial multi-item working memory studies (15,86–88). At high loads, merging and loss of memories occurred in the network, mimicking forgetting errors for multi-item working memory (37,40,41,89) (example load-4 trial in Fig. 7a). When each area reached its capacity, preferentially contralateral items were stored, similar to the simpler model (Fig. 6). For this reason, when analyzing the report accuracy (see Methods) in load-4 trials, we observed that memory errors increased the more non-target items were displayed in the target hemifield (Fig. 7b), replicating the bilateral field advantage (15).

**Figure 7:**
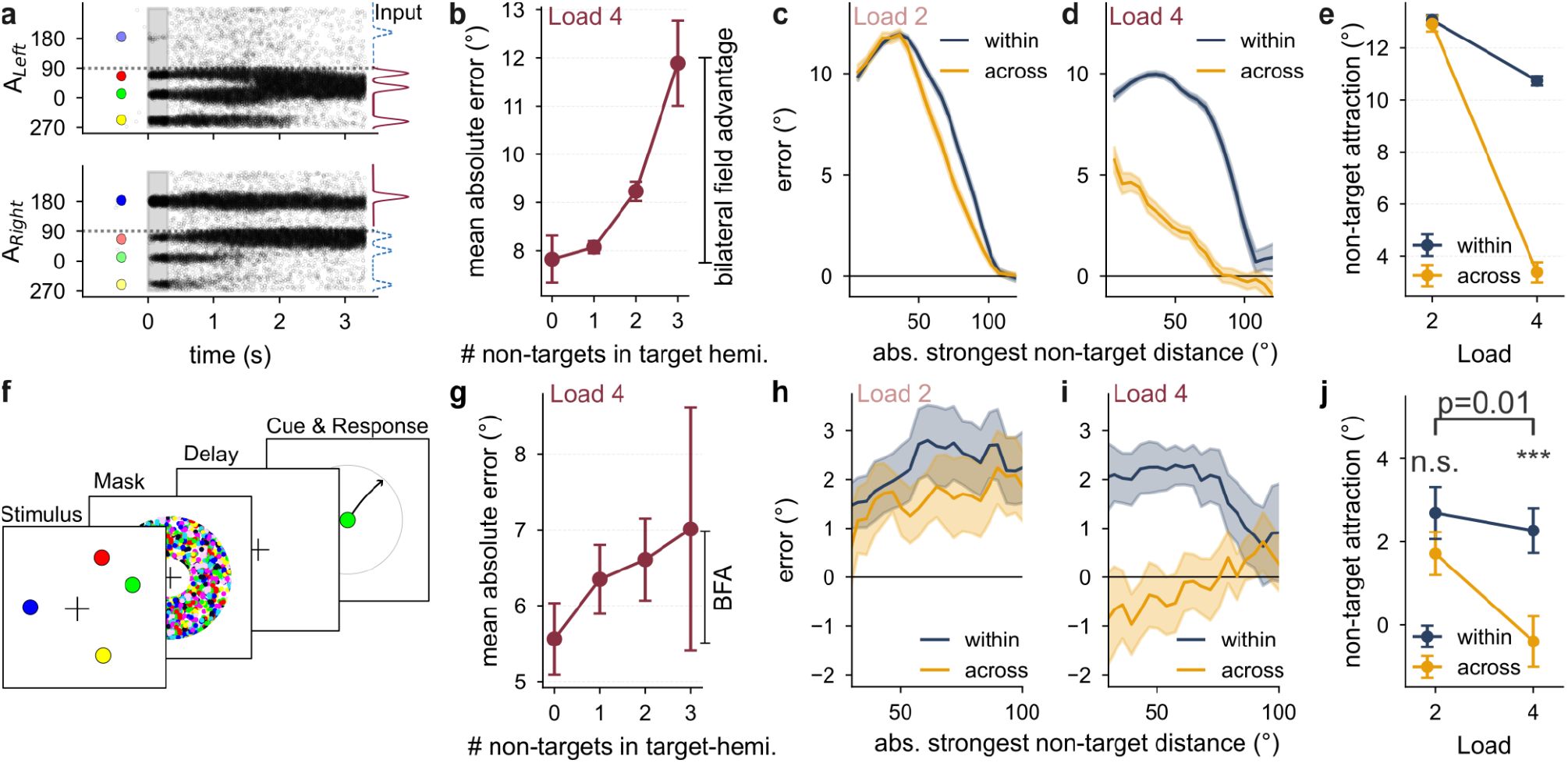
Human working memory shows load-dependent changes of attraction towards non-targets consistent with model predictions. a-e) Higher-capacity Model (n=6,000 trials): a) Example simulation for four presented targets (colored dots for illustration only). Items can merge (red and green, Area_Left_), fade (e.g. yellow, Area_Left_) or be maintained (blue, red, Area_Right_). b) Mean absolute error of simulations increases the more non-targets are presented in the target hemifield for load-4 trials, akin to the bilateral field advantage (BFA). c) Errors in relation to the absolute relative distance of the target and non-target for load-2 trials when the non-target was presented within the target-hemifield (within) or the opposite hemifield (across). d) Same as panel c for load-4 trials when using the strongest non-target (estimated by a DoG1-fit, see Methods). e) Non-target attraction (DoG1-fit) of panel c, d (estimated by DoG1 fits) comparing within and across hemifield attraction for the different loads. f-j) Human experiments (n = 18 subjects): f) Task design (90,91): Stimuli are simultaneously presented, masked and the color-cued location is reported after a delay. g) Same as panel b for the data (Mixed effect model across participants ‘absolute_error ∼ hemifieldLoad’, two-sided; β = 0.58 °/item, 95% CI [0.18, 0.99], t(2056) = 2.83, p = 4.7×10^-3^, N = 4942 trials across 18 subjects). h, i) Same as panel c, d for the data, showing within vs across hemifield non-target attraction for load-2 and load-4 trials, respectively. j) Summary of panel h, i showing non-target attraction (DoG1 fit) for the within and across hemifield conditions. Higher load increased the difference of hemifield-specific non-target attraction (paired two-sided t-test of within-across DoG1 fits: t(17) = −2.87, p = 0.011, mean = −1.69, 95% CI [−2.94,-0.45]), with no hemifield-specific difference for load-2 trials (t(17) = 1.82, p = 0.086, mean = 0.98, 95% CI [−0.15,2.1]) but a significant difference for load-4 trials (t(17) = 4.04, p = 8.48×10^-4^, mean = 2.67, 95% CI [1.27,4.06]). Error bars represent ±1 s.e.m.

Arguably, increased errors when more items are placed in the same hemifield as the target partly reflect shorter distances between items, which are known to increase memory interference (37,89). In order to separate the role of distance-based interference from that of specialized hemifield storage in high load trials, we designed an analysis conditioned on inter-item distance. We computed memory interference as the response error relative to the location of the most interfering non-target item (see Methods). Importantly, this non-target attraction can be analyzed separately for non-targets within the target-hemifield or across hemifields.

We found that when the network operated below its capacity limit (load 2), interference by non-targets within and across hemifields was comparable (Fig. 7c), reflecting the fact that both items were often redundantly stored in both areas (example trial in Supplementary Fig. 10b). In contrast, when the network operated above capacity (load 4), mostly the stronger, contralateral stimuli were successfully encoded in each area (example trial in Supplementary Fig. 10c), resulting in reduced interference by items presented across hemifields relative to those presented within the target-hemifield (Fig. 7 d, e), even when conditioning on inter-item distance.

In our computational model, this load-dependent change reflects a switch in storage strategy from redundant to specialized hemispheric representations. Crucially, this can be readily tested behaviorally. To this end, we pooled data across two publicly available human behavioral data sets (90,91). 18 participants performed a spatial multi-item working memory task from which we analyzed trials in which two or four colored circles were briefly presented at different spatial locations. Following a colored mask and a delay period, the target was cued by a colored circle at the fixation point, and participants responded by moving the item to the remembered spatial location (Fig. 7f). Precision and attraction towards non-targets was comparable across the data sets, allowing us to pool across them (Supplementary Fig. 11 a,b,e,f). For fixed memory load at four items, memory errors increased with the number of non-target items in the target hemifield, revealing a bilateral field advantage (Fig. 7g).

We next examined the distance-dependent memory interference. As predicted by the model, the low load condition (load 2) showed similar distance-dependent attraction towards non-target items presented within or across hemifields (Fig. 7h, j). However, at high load (load 4), attraction towards non-targets presented across hemifields was almost entirely abolished (Fig. 7i, j). Indeed, we found a significant triple-interaction between the non-target attraction fits of within and across conditions with increasing load (p=0.01, Fig. 7j, hyperparameter crossvalidation in Supplementary Fig. 11 c,d,g,h, see Methods). This load-dependent change in memory interference (Fig. 7j) mirrors our computational model predictions (Fig. 7e), indicating that redundancy at low task demands is flexibly replaced by specialized hemispheric storage at high memory loads.

Taken together, our results demonstrate that redundant, weakly connected hemispheres with lateralized inputs flexibly optimize working memory performance based on task demands. In simple below-capacity tasks, the redundancy allows for increased precision, while in more demanding above-capacity tasks independent stimulus activation allows for increased capacity through hemispheric specialization.

## Discussion

We explored the prefrontal architecture supporting interhemispheric spatial working memory. While mnemonic spatial representations exhibited a contralateral bias in monkey PFC (Fig. 2g,h), each hemisphere predicted trial-to-trial behavioral errors equally well (Fig. 3b,c) and had weakly correlated encoding errors (Fig. 3e,f). These findings align with a network model where each hemisphere operates as a separate attractor network, with tuned but weak interhemispheric connectivity (Fig. 4a-e). This architecture allows for shared memory diffusion (Fig. 4e) as well as private history effects such as reactivations and memory drift (Fig. 4f-k), consistent with the experimental data (Fig. 5). Finally, we demonstrate that this architecture can represent working memory items redundantly or independently, depending on working memory demands (Fig. 6) and confirm model-derived predictions in a human behavioral data set (Fig. 7).

Importantly, this study addresses a pending debate about interhemispheric working memory organization (5–7). On the one hand, multi-item working memory studies suggest largely independent resources in each prefrontal hemisphere, supporting a lateralized organization (Fig. 1a; reviewed in (7)). On the other hand, optogenetic studies in mice indicate a redundant organization, since the disruption of both hemispheres is necessary for memories to be irrevocably disrupted (5,23,24). Our model reconciles these views by proposing a redundant, weakly connected network with lateralized inputs. When memory demands are low, both hemispheres are redundantly activated, improving memory precision and robustness. When demands are high, competition for lateralized resources promotes hemisphere-specific encoding and increases working memory capacity. Importantly, this flexibility emerges from input lateralization alone, without requiring task-dependent changes in connectivity, as proposed in previous studies (19,23,26,92). Taken together, this network flexibly adapts to varying cognitive demands with a fixed, parsimonious architecture.

This architecture seems to strike a functional trade-off between redundancy and specialization reminiscent of strategies used when storing data in multiple hard drives. When storage space allows, maintaining back-ups reduces the probability of information loss. As storage space becomes limited, prioritizing unique content over duplicates becomes more efficient. The prefrontal hemispheres seem to implement this principle by combining a redundant structure with hemisphere-specific inputs, prioritizing either robustness or capacity depending on task demands.

We used decoding analysis to focus on interhemispheric interactions related to selective memory encoding, and found that within-region correlations along this neural space are stronger than cross-regional ones (Fig 3e). This is in contrast with interhemispheric activity in neural subspaces independent of selective memory encoding, which show much stronger correlations across hemispheres (54), suggesting that arousal signals are more globally shared than working memory related activity.

Our observation of correlated neural errors between the hemispheres led us to infer the existence of weak, tuned interhemispheric connections, consistent with observed interhemispheric noise correlations in these data (see Fig. S2 in (54)). While one may argue that these correlations rather reflect common input, such input would need to induce consistent rotations of mnemonic representations in both hemispheres to explain the alignment of small memory imprecisions. Uninstructed eye movements, the most plausible confound, did not alter the effect, though correlated noise from upstream areas could provide an alternative explanation. However, even if an unidentified confound contributed, our model predictions beyond shared memory diffusion should remain largely intact, since the unconnected and weakly connected models from Fig. 4 show no qualitative differences.

While our analyses likely support weak interhemispheric connections, other studies suggest stronger coupling. Specifically, neural selectivity in the mouse ALM can reemerge from the opposite hemisphere after unilateral optogenetic disruption (23,24,93). This discrepancy may reflect task-dependent increases in background input that facilitate faster recovery during active engagement (93) or differences in targeted brain regions. Notably, the optogenetic studies were conducted in the mouse premotor cortex (23,24,26,93), an area known to have stronger callosal connections than PFC (94,95). These distinctions highlight the need to investigate how memory lateralization varies across the cortical hierarchy, adding a lateral dimension underexplored in studies that emphasize the role of distributed brain processing in working memory maintenance (40,52,53,96,97), serial dependence (98), and multi-item working memory (9,15,21).

Finally, our findings can have implications for several neuropsychiatric disorders, such as schizophrenia and autism, which show a reduction in callosal connectivity (99,100). The observed deficits in working memory performance in these disorders (34,101,102) should be researched in light of the changes in interhemispheric connectivity.

In conclusion, our study bridges paradoxical findings from single- and multi-item working memory studies by providing a single functional architecture underlying prefrontal hemispheric organization. The proposed redundant, weakly connected architecture accounts for memory effects across task complexities and timescales, encompassing memory precision, history effects, and the bilateral field advantage in multi-item working memory.

## Methods

### Behavioral task and recordings

All experimental procedures were approved by the Institutional Animal Care and Use Committees of Carnegie Mellon University and the University of Pittsburgh, and were in accordance with the United States National Research Council’s Guide for the Care and Use of Laboratory Animals. Three male rhesus macaque monkeys (Macaca mulatta) were used in this study. One animal was sourced from Tulane National Primate Research Center (Monkey Wa) and two animals were from Covance (Monkey Pe and Monkey Sa). They were 10 (Monkey Pe), 7 (Monkey Wa), and 7 (Monkey Sa) years old at the time of data collection. Animals performed an oculo-motor delayed response task (103), where they had to fixate on the center of the screen while a peripheral stimulus was shown for 420ms in one of 4, 8 or 16 different locations (7 (3,453 trials), 12 (5,666 trials), 22 (9,065 trials) sessions, respectively). After a delay period the monkeys made a saccade to the remembered stimulus location to report their response and gain a water reward for correct trials. These data were collected as part of a separate study (54,55), which focused on the link between shared interhemispheric variability and arousal signals, independently of selective memory encoding. Instead, we analyze here neural activity and behavior related to the selective encoding of spatial memories, irrespective of global non-selective co-variations. The details of the experiments are briefly summarized here.

We excluded all trials with fixation breaks and only utilized rewarded trials (29,459 trials out of 64,504 trials were rewarded (45.67% rewarded trials)). Furthermore, we excluded trials with errors larger than 22.5° (291 trials excluded, ∼1%) and only analyzed consecutive correct trials, leaving a total of 18,184 trials (61.72% of all rewarded trials). Consecutive correct trials were selected to avoid potential confounds arising from fixation breaks in the preceding trial, which accounted for the majority of excluded trials, and to prevent ambiguity introduced by incorrect responses on the preceding trial.

Delay lengths were randomly varied between 1.5s and 3s, where each session uniformly covered a 0.5s interval of delay lengths (except session Sa1, which covered 1.5s) and each monkey had a different range of delay lengths (Sa: 1.5s - 3s, Pe: 1.5s - 2.6s, Wa: 1.5s - 2.4s). Between trials, a mostly self-timed inter-trial interval of 0.9s - 2.2s separated individual trials. Fixation was maintained within an invisible window around the fixation point (Monkey Sa: 2.4 or 2.9, Monkey Pe: 2.4, Monkey Wa: 2.8 degrees radius) throughout the entire trial until response. However, we here denote the fixation time as the time from fixation onset until stimulus presentation. Eye Tracking was performed with an EyeLink 1000 eye tracker system with a sampling rate of 1 kHz.

All three monkeys were implanted with a 96-channel multi-electrode recording device (Utah array, Blackrock Neurotech) in each hemisphere of the PFC anterior to the arcuate sulcus and dorsal to the medial sulcus (area 8Ar), which were used to simultaneously record activity during the task. The recordings led to a total number of 5708 recorded multi-units across both hemispheres for all animals and 41 sessions (Sa: 137.9±16.4 units/session, Pe: 157.9±15.4 units/session, Wa: 108.4±10.8 units/session). We excluded one session from monkey Pe from most neural analyses, due to a shorter stimulus being used. We excluded multi-units with firing rates lower than 1 Hz for the analyses.

Monkey Pe tended to have a smaller amount of selective neurons in each session (average of 45 selective units per session) than the other monkeys (121 (monkey Sa) and 78 (monkey Wa) selective units per session), likely due to sampling differences of the implanted array.

### Human data: Multi-item working memory study

We combined two publicly available data sets of versions of the same task (90,91). A total of 18 healthy participants (n=10 in Data1; n=8 in Data2; 10 females, 8 males, 21-37 years old) performed a total of 5066 trials. The experiment is described in detail in the original studies and we only note here that between 1-8 items were presented across both data sets (1, 2, 4 items in Data1; 1, 2, 4, 8 items in Data2), of which we only analyzed trials with a set size larger than one item. Subjects fixated in the center of the screen while different colored stimuli were presented across a circular space. A colored mask was presented after stimulus presentation, which was followed by a variable delay (Data1 variable: 0.5s, 1s, 2s or 4s; Data2: fixed 1s). Subjects were then cued by a colored item appearing at the fixation cross and responded with the location of the cued item. Subjects responded either with their eyes (Data1) or hands (Data2). We excluded trials with errors larger than 50° (Data1: excluded 97 out of 2560 trials; Data2: excluded 156 out of 2635 trials). Performance for both data sets was comparable so we pooled across experiments (Supplementary Fig. 11).

### Statistics & Reproducibility

All analyses were performed in Python (version 3.6.13). Statistical tests (t-test) were performed using the SciPy (version 1.5.2, https://scipy.org) statistics module, linear models (OLS) were performed using the Statsmodels (version 0.12.2, https://www.statsmodels.org/) module. Circular correlations were performed using the Pingouin package (version 0.4.0, https://pingouin-stats.org/). Gaussian Mixture Models and crossvalidations were performed using the Scikit-learn library (version 0.24.2, https://scikit-learn.org/). Attractor network simulations were performed using Brian2 (version 2.4.2, (104)). Sample sizes and p-values are directly reported in the corresponding figure caption. In studies involving non-human primates, by convention in the field a result in one animal needs to be replicated in a second for the finding to be considered robust. We used 3 animals in this study, as collected for the original publication (54). In addition, most of our statistical measures are calculated across sessions and not across animals, indicating that the number of sessions is the relevant sample size. For human studies the original studies estimated sample sizes (90,91). We combined two studies for more reliable estimates. Averaged data shown in figures are averaged across non-human primate sessions (n = 40), simulations or human participants (n = 18), noted in each caption. Detailed descriptions of plotted data are provided in each caption. If data were excluded it is mentioned in each caption.

## Behavioral data analysis

### Precision analysis

Each trial’s saccadic endpoint response was parameterized in polar coordinates taking the fixation point as origin and only the angular dimension θ_r_ was considered for further analysis. In each trial, the angular error (θ_*e*_) was computed as the circular distance between the stimulus angle (θ_*s*_) and the response (θ_*r*_). We subtracted the mean of θ_*e*_ in each target location for each subject, session and delay to account for systematic biases (except for the analyses shown in Fig. 1d,e). For each monkey, precision was estimated by fitting an ordinary least squares (OLS) model on the absolute

response error with the delay (abs(θ_*e*_) ∼ delay) across sessions using the statsmodels.formula.api Python package.

### Serial dependence plots

To visualize serial dependence (Fig. 1f), we plotted the current trial’s error 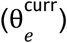 against the circular distance between the previous and the current stimulus location 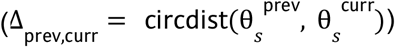.

### Serial dependence strength fit

To measure the strength of serial dependence β, both in behavior and neural analyses, we fit behavioral or decoder errors to Δ_*prev,curr*_ using an OLS model: θ_e_ ∼ β *F*(Δ_*prev,curr*_). The typical shape of serial dependence curve *F*(*x*) was parameterized using a negative first-order normalized derivative of Gaussian (DoG1) (34):

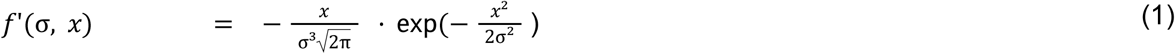

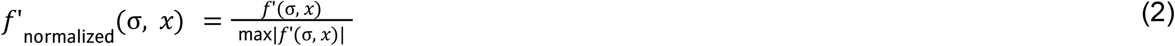

For each monkey, we determined *F*(*x*) = *f*’_normalized_ (σ _opt_, *x*), where σ_opt_ was obtained for each monkey by minimizing the model’s Bayesian information criterion (BIC) within a range of σ values (σ = [0. 3, 3. 0, 0. 05] rad) for all trials from that monkey (Supplementary Fig. 7, a-c for behavior, d-f for neural history drift, g-i for neural reactivation drift).

### Determining the most-interfering non-target

In the multi-item spatial working memory task (Fig. 7f), we determined the most-interfering non-target item, i.e. which of the presented non-target memoranda would maximally attract the target memory. To do so, we first optimized the DoG1 for each data set for each set size separately. For a range of σ (range from 0.1 to 2.0 radians in 0.05 steps), we determined in each trial which of the presented non-target items would theoretically maximally attract the target. We then fit a DoG1 with the same sigma on the response errors over the relative distance of the target to the theoretically most-interfering non-target. In this manner, we determined an optimal hyperparameter for each data set and each load based on the BIC of the model fits (Supplementary Fig. 11 c,d,g,h). For Data1 we obtained σ^*D*1^ _opt,load2_ = 1. 4 for the load-2 condition and σ^*D*1^ _opt,load4_ = 0. 8 for the load-4 condition. For Data2 we obtained σ^*D*1^ _opt,load2_ = 1. 1 for the load-2 condition and σ^*D*1^ _opt,load4_ = 0.7 for the load-4 condition. We recomputed the relative location of the most-interfering non-target for σ_opt_ for each load and used this to evaluate distance-conditioned memory interference.

## Neural data analysis

### Memory fields

To compute the preferred location of each multi-unit during memory maintenance (memory field (MF)), we obtained its mean firing rate in each trial *r*_*i*_ (spike count between stimulus offset and go-cue divided by delay duration) for each stimulus location (θ_*s,i*_) and computed the population vector (31):

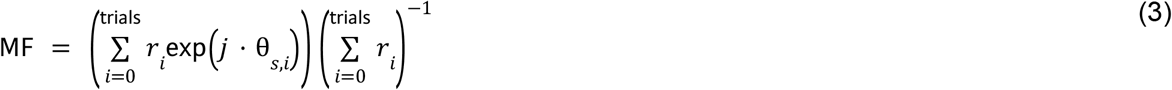

The preferred orientation of each multi-unit θ_pref_ is given by the angle of the complex-valued MF and the tuning strength by the absolute value of the MF.

To determine if a multi-unit’s MF tuning was significantly above chance, we computed the MF for 1000 random shuffles of stimulus locations. A multi-unit’s MF was classified as significant when its tuning strength exceeded 97.5% of the tuning strengths of the shuffles.

### Visualizing PFC bumps

To visualize the representation of the stimulus in PFC activity (Fig. 2b,c), both in individual trials and as an average over trials, we displayed the firing rate of each multi-unit over time, where the multi-units were sorted by their θ_pref_. To avoid the circularity of generating a bump during the delay solely by sorting by θ_pref_, we estimated θ_pref_ in trials not used for display using a 5-fold cross validation for trial average, and a leave-one-out cross validation for single trial visualization. To correct for heterogeneous baseline firing rates of each multi-unit, we subtracted the average firing rate (across time and trials) from each multi-unit’s activity.

Time-resolved population representations were obtained by averaging both along the time and the neuron axes. Along the time axes, we smoothed firing rates with a sliding average window (step size 10ms; window size 100ms for trial averages, and 200ms for single trials). Along the neural axis, we also performed a moving average smoothing (bin width W = 2π*N*^−1^, with N the number of different target stimuli used in the session; step size 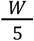).

Finally, prior to averaging representations across trials we re-centered the multi-unit’s relative to the stimulus angle in each trial, so that multi-units preferring each trial’s stimulus location were placed at the axis origin.

### Population decoding of stimulus location

We used linear regression to decode stimulus location from population activity in the trial. For each unit and trial, spikes were binned into independent 200ms time bins aligned to fixation or response onset. Due to variable delay lengths, the fixation- and response-aligned delay activity share a partial overlap since a section of the shortest delay duration (for each session) from stimulus start or delay-end is taken into account.

All decoders were trained using a 5-fold stimulus-stratified cross validation approach. In the training set (80% of the trials), at each time *t* the complex weights β_*t*_ of the neural activity regressors *X*_*t*_ were estimated to reproduce the complex stimulus location *y*_*s*_ by least-squares estimation:

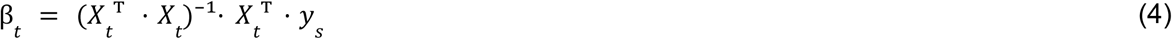

where both β_*t*_ and *y*_*s*_ are complex vectors, describing the coordinates (*a* + *ib*) of the trained receptive field and stimulus locations, respectively.

The computed weights were then used to predict the stimulus location 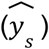 based on the neural activity in the previously left out trials (test set, 20% of trials):

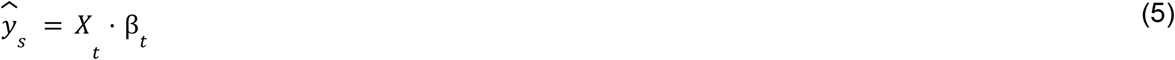

As a metric for the decoding strength of the trained decoder, we then evaluated the mean squared error (MSE) of the angular predictions 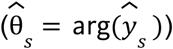 against the angular stimulus locations (θ_*s*_ = arg(*y*_*s*_)):

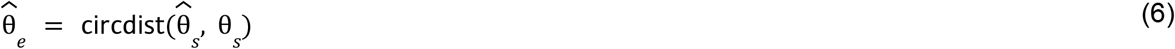

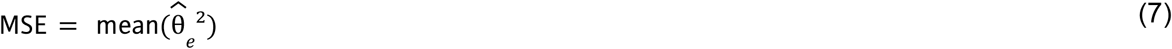

To account for the different number of stimuli in each session and any potential unbalanced trial distribution, we also tested the previously trained model repeatedly (50 times) against randomly shuffled θ_*s*_. To compute a decoding score, we z-scored the MSE against the shuffled MSEs (MSE_shuffle_) and provided the output as a measure of standard deviation σ. The decoding score is thus obtained as:

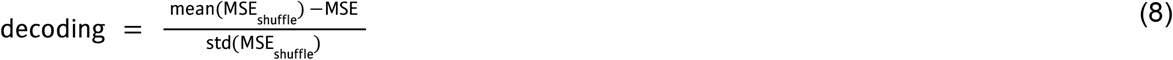

where we changed the sign of the score to facilitate interpretability, so that high values correspond to high accuracy.

#### Cross-temporal decoders

We trained a linear regression to fit the stimulus location as a linear combination of the neural activity in each time point using a stratified 5-fold cross validation. For each trained time point, we tested the model on all time points of the left-out trials. The cross-temporal decoder matrices shown in Fig. 2e-g only shows fixation-aligned activity for simplicity.

#### Single-trial decoders

While the cross-temporal decoders were used to gain a general impression of the data, the single-trial decoders were meant to give a precise prediction for individual trials. We focused on two different decoders: A Same-Time Decoder for estimates based on each time step (train, test in same time point; diagonal of cross-temporal decoder) and a Delay Decoder which only contains the memory code (train on average delay activity, test in each time point). The Delay Decoder allowed for the detection of memory reactivations at the next trial’s fixation onset. As shown before (32,50), the Same-Time Decoder included response-related activity. For single-trial decoders, we performed a 20-times repeated 5-fold cross validation. We performed the repeats since single trials were noisy and averaging across repeats led to more stable predictions.

### Decoder-behavior error correlations

We computed trial-by-trial correlations between neural decoder prediction errors 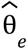 and behavioral response errors θ_*e*_ to view how prefrontal population codes reflected the small clockwise and counter-clockwise response errors of the monkeys. We either trained one decoder on all multi-units across hemispheres (combined analysis, Fig. 3a) or one per hemisphere (independent analysis, left vs. right hemisphere, Fig. 3b,c). The obtained prediction errors (combined, left, right) were each circularly correlated (circ_corrcc from pingouin package (version 0.4.0) (104)) with the behavioral response errors in each session in each time step and averaged across sessions (error bars taken across sessions). In the independent analysis, we performed the correlations either on only ipsilateral vs. contralateral trials relative to the respective hemisphere, or for the left and right hemisphere separately on all trials, to compare which hemisphere predicts behavior.

### Decoder-decoder error correlations

We assessed task-related coupling between simultaneously recorded neural populations of the two hemispheres by estimating the trial-by-trial error correlations between predictions of the respective decoders (Fig. 3e,f). We compared within hemisphere to across hemisphere error correlations. To do so, we repeatedly (5 times) trained four independent decoders on a random half of units from each hemisphere (left_A,_ left_B_, right_A._ right_B_) respectively. The obtained decoder errors 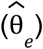 were then circularly correlated between the different decoders, either within a hemisphere (left_A_ vs. left_B_, right_A_ vs. right_B_) or across hemispheres (left_A/B_ vs. right_A/B_, respectively) at each time point for each repeat and session. The correlations were averaged across the five repeats and afterwards across sessions (error bars are across sessions).

### Uninstructed eye-movement controls

We used a Same-Time Decoder to decode the stimulus location from uninstructed eye movements (Supplementary Fig. 6a). We computed the decoder as mentioned previously, but we replaced the multi-unit activity by the complex gaze location (see section Population decoding of stimulus location). Since the acquisition rate of the eye tracker and the used, preprocessed acquisition rate of the electrophysiology (original acquisition rate 30kHz, downsampled to 1 kHz) were the same, we binned the eye locations in the same manner as the neural data.

We also correlated the gaze location with the stimulus location (Supplementary Fig. 6b, angle) and the gaze error with the performed response error (Supplementary Fig. 6b, error), to understand if gaze location could correlate performance errors across hemispheres. Since gaze and later response errors were not correlated we excluded this confound.

Additionally, we tested if removing the information of the complex gaze location from the neural decoder had an impact on error correlations (Supplementary Fig. 6c,d). To do so, we added the gaze location as an additional regressor to the neural activity in each of the four sub-hemispheric decoders (complex gaze location is treated like an additional multi-unit). However, we computed the error predictions by testing on a partial model, where gaze information was discarded. This gave us neural predictions that did not overlap with stimulus encoding through uninstructed eye movements. The gaze-excluded predictions were correlated within and across sub-hemispheres (left_A,_ left_B_, right_A._ right_B_) as described above (Supplementary Fig. 6c, d).

Taken together, these analyses control for uninstructed eye movements as a potential source of common input, but other uninstructed movements also elicit shared neural variability across brain regions although less in monkeys (105) than in mice (106). However, the shared neural information underlying correlations in memory imprecisions is highly specific and can only be conceivably induced by movements that cause a consistent rotation of mnemonic representations across hemispheres. We therefore argue that eye gaze is the main confound to address, as it has been shown to change spatial encoding during the delay period (75,79,107). Since removing eye-confounds did not change the correlations, we argue for a task-related communication subspace (108) underlying information transfer between the prefrontal hemispheres, likely realized through tuned interhemispheric connections anatomically supported by the corpus callosum.

### Neural history drift

To estimate the attraction (or repulsion) of the neural population code in the current trial’s delay period towards (or away from) the previous trial stimulus location, we plotted the single-trial Sametime Decoder prediction errors 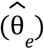 against the distance of the previous and current stimulus (Δ_prev,curr_) (Fig. 5a). We quantified this history drift by fitting a negative first-order normalized derivative of Gaussian (DoG1) to this curve, as for the behavior (see section Serial dependence strength). The hyperparameter σ_opt_ was chosen based on the model BIC (see section Serial dependence strength; Supplementary Fig. 7, d-f, Sa: 1.35 rad, Wa: 1.75 rad, Pe: 3.0 rad). We applied this fitting procedure to 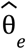 in each time step (200ms) through the stimulus and delay periods (Fig. 5b).

To account for auto-correlation between sequential stimuli (consecutive trials were less likely to occur in the same location) we computed the decoder errors as the circular distance between 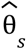 and a shuffle control 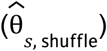. We obtained the shuffled predictions 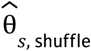 by shuffling the neural activity of trials with the same current stimulus location randomly (50 repeats) and multiplied them with the decoder weights. This corrected for uneven sequential stimuli distribution, while retaining the drift towards a specific previous stimulus location (as only trials with the same current stimulus are shuffled). This shuffle was only performed for completeness to extrapolate the drift to the previous trial. It did not affect the drift of the current trials shown in Fig. 5b.

### Reactivations

Fig. 2h shows that reactivations occur immediately before stimulus presentation. We therefore estimated the location and strength of reactivations in individual trials using a Single-trial Delay-Decoder on neural activity averaged over the two time bins in the time window starting from fixation start until the current stimulus start. The angle of this decoder’s prediction provides the reactivation location, while its absolute value reflects the reactivation strength.

### Reactivation drift

The reactivation drift was computed in a similar manner to the neural history drift but instead of considering the relative location of the previous trial’s stimulus, we considered the relative distance of the current stimulus to the previous reactivation location (Δ_react,curr_). Therefore both components of the reactivation drift, the relative reactivation locations and the delay prediction errors, depended on the neural predictions from either hemisphere. We could therefore compute the drift of the population activity during the delay towards the reactivation either within the same hemisphere or across hemisphere (Fig. 5d).

### Gaussian Mixture Model

To analyze if reactivations co-occur in the two hemispheres we applied the Gaussian mixture model (GMM) from scikit-learn to the scatter plot of trial-by-trial reactivation decoder strengths from the right and left PFC hemispheres (Fig. 5e). We decided on modeling four classes as the hypothesis was for the model to find a class for each: null, single_left_, single_right_ and dual reactivations. We determined the covariance constraint by comparing the BIC of the different models (best fit: diagonal covariance) and repeated the GMM fit 250 times with different initial parameters. The fact that dual reactivations (a class along the diagonal indicating strong decoder strength for both hemispheres) did not appear in any of our fits of reactivation strengths (Fig. 5e) but were instead systematically observed if instead of reactivations we considered the strength of delay-period codes (Fig. 5f) indicated that reactivations occurred privately in each hemisphere.

### Bump-attractor model

We adapted the previously proposed bump attractor model implementation from (32) based on (109) in Brian2 (110). We simulated 1024 excitatory and 256 inhibitory leaky-integrate-and-fire spiking neurons for each area representing the left and right hemispheres. Within each area, all neurons were densely connected (all-to-all connectivity) with AMPAR, NMDAR and GABA_A_R synaptic dynamics (decay time constants τ_AMPA_ = 2ms, τ_NMDA_ = 100ms, τ_GABA_ = 10ms). The strength of excitation between excitatory neurons in an area depended on the distance between their respective receptive fields through a Gaussian connectivity profile (std 14.4 deg). All other within-area connections were untuned.

Most within-area parameters were kept as in the original implementation (32), but we changed *J* _*EE*_ ^+^ = 1. 73, η_poisson_ = 0. 925Hz.

#### Short-term plasticity

The model also incorporated short-term synaptic plasticity in the excitatory-to-excitatory connections within each area (32), following the dynamics of facilitation (*u*) and depression (*x*) proposed by (80):

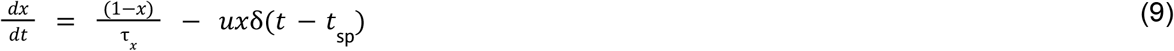

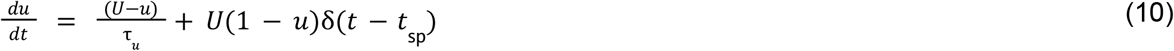

where *t*_sp_ marks the spike times and δ(*t*) is the Dirac-Delta function. The constants were chosen as τ_*x*_ = 0. 25s, τ_*u*_ = 1. 5s and *U* = 0. 15, and variables were initialized as *u*_0_ = 0. 08, *x*_0_ = 0. 9. Synaptic weights between excitatory neurons were multiplied by *u* · *x*. The addition of short-term plasticity to the bump-attractor model allows for activity-silent storage of previous spiking activity and therefore allows for serial dependence through the inter-trial interval as previously shown (32).

We also implemented a perceptual repulsion to the previous stimulus by shifting the stimulus input a maximum of 0.75° away from the previous stimulus through a DoG1 (σ = 0. 5 rad) of the distance between the previous and current item Δ_prev,curr_, similar to (34,35).

#### Across-area connectivity

We connected the two area networks for the left and the right hemispheres via excitatory connections either in a tuned (scaled version of the within-area excitatory-to-excitatory connectivity) or random manner. Tuning was defined solely based on the distance between preferred locations of pre- and post-synaptic neurons across the two area networks. An alternative tuning function may additionally depend on the preferred location of the presynaptic neuron, which would allow modeling midline connectivity inhomogeneities. However, our analysis of midline versus peripheral error correlations did not show any evidence for midline specificity (Supplementary Fig. 6e), and we therefore did not investigate this connectivity structure further. For simplicity, across-area connections were only mediated by AMPAR excitatory-to-excitatory (EE, tuned) and excitatory-to-inhibitory (EI, untuned) synaptic dynamics as in (36). To stay within a similar parameter regime when varying across-area connectivity, we compensated the across-area connectivity by concomitant decreases in within-area EE and EI AMPAR connectivity. We simulated a total of N = 60,000 trials and varied the tuned across-area connectivity between 0.2% and 1% of the within-area baseline model strengths (in 0.2% steps, one randomly connected condition, resulting in ∼8500 simulations per condition).

#### Stimulation paradigm

An example simulation can be seen in Figure 4b. A positive current (0. 3nA) of a Gaussian shape (σ = π/2 rad) was selectively applied to the neurons centered around the chosen stimulus location for 500ms. Following the stimulus extinction, circuit reverberation supported tuned persistent activity that lasted through a 3s delay before a negative, non-specific input to all neurons (− 0. 6nA, 500ms) disrupted the delay activity and reset a low-rate baseline state. The network was left unperturbed and mostly silent for an inter-trial interval of 1500ms, after which we applied a reactivation signal through a weak, positive, non-specific input (0. 036nA, 500ms). This input reactivated the previously silent code in 38% of trials and was turned off after a period of 300ms with a negative stimulus (− 0. 3nA, 500ms). After another 1200ms, the second stimulus input was applied at a random location with the same characteristics as the first stimulus input. However, its stimulus location was shifted slightly away from the previous stimulus to simulate sensory adaptation from upstream areas (see section Short-term plasticity). This stimulus also elicited persistent activity, which was maintained during another 3s delay, after which the simulation was terminated.

#### Model behavior

The behavioral readout (θ_bump_) from each area of the model was determined by computing the circular mean of the excitatory neurons’ preferred direction (θ_*n*_) weighted by their firing rates (*r*_*n*_):

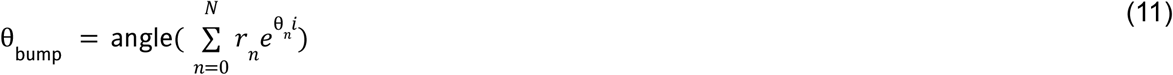

with *N* being the number of excitatory neurons. The equivalent of a behavioral response of the model simulation was determined using the firing rate average in the last 250ms of each delay. We also computed neural decoder readouts using the same approach but considering *r*_*n*_ in 200ms time windows (in 100ms steps) throughout the trial. The reactivation location and strength was computed using the same decoding approach with *r*_*n*_ in the 250ms following the reactivation stimulus end.

#### Multi-item simulations

For the simulations in Figure 6, the same model as in the single-item working memory simulations (Fig. 4) was used with an across-area connectivity of 0.2% (which fits the data in Fig. 5) and simulated for n=1500 trials. The only change to the network was the introduction of a slight stimulus input lateralization (84): We assigned each area to a hemisphere (Area_1_ = left, Area_2_ = right) and varied the stimulus strength according to the target’s hemifield (90-270° = left; 0-90° or 270-360° = right). When stimuli were presented contralaterally to a hemisphere, the stimulus strength was set to 0. 3nA, as before, but if the stimuli were presented ipsilaterally the stimulus strength was reduced to 0. 21nA (70% of contralateral strength). The results remained qualitatively true for varying the ipsilateral stimulus strength in a reasonable range.

Both networks were simultaneously stimulated with two stimuli, where one stimulus was always induced at 45° (right hemifield), while the other stimulus was randomly presented in 110° distance to the first stimulus, placing it at 155° (left hemifield, across hemifield to the first) or 295° (right hemifield, within hemifield to the first). The time course of the stimulation protocol remained the same as in the single-item task.

For this proof-of-concept we reused the model from the previous analysis, which has a default capacity of one item per ring. This led to a system-wide capacity of two-items since we have two attractor networks (Area1, Area2). However, fine-tuning the model with stronger and more tuned excitation (41) or inhibition (40) can increase the working memory capacity of each bump-attractor model, which would allow modeling of higher capacities.

To read out two simulated behavioral responses from each model simulation, we divided the network into three equally sized bins (0-120°, 120-240°, 240-360°) and decoded from the neural activity in the two bins where stimuli had been presented. This either led to a precise readout if the bump for the corresponding stimulus had been maintained or to a random location readout in case the memory was not retained. This allowed us to determine the network capacity, meaning how many items were retained in each trial (Fig. 6e).

The memory precision (Fig. 6f) was computed by determining which stimulus was retained in each area and computing the response error to the remembered stimulus. We compared simulations where one item was remembered (the same item in both areas) to simulations where both items were remembered (one item per area).

#### Higher-capacity model

We changed three parameters of the network to allow for a higher within-network capacity and simulated it for n=6,000 trials. We reduced the stimulus width from 60° to 50°, reduced the excitation weight profile (σ_weight_) from 14.4° to 6.1° and increased *J* _*EE*_ ^+^ from 1.73 to 2.28.

We presented the network with either two (low load condition) or four stimuli (high load condition), where all stimuli had to have a minimal distance of 30° from each other, as in the Multi-item data set (90,91). Apart from adding more stimuli, the simulations remained the same.

To extract response errors from our simulations, we randomly selected an item as the target stimulus. We then formed a bin around the target location (width = ±30° around the target) and decoded the location of the corresponding response with a population vector decoder based on 200ms late-delay spike counts of neurons contained in the corresponding subsection. If the absolute value of the complex population vector was below a threshold of 0.5 a random location was chosen as the response. This cut-off could be well established due to a bimodal distribution of population vectors for trials with and without memory representations for an item. As in the data, we excluded simulations with errors larger than 50°. We estimated the most interfering non-target item by optimizing the fit of a DoG1 function akin to the data (see section Determining the most-interfering non-target) on all trials, obtaining σ_opt,model_ = 0.65. We then computed which non-target maximized the fit of the DoG1, obtaining the theoretically most-interfering non-target item and used this to evaluate distance-conditioned memory interference in the model.

## Data availability

The non-human primate data (54) analyzed in this study are available at the KiltHub database from Carnegie Mellon University (55). The human data (90,91) analyzed in this study have been retrieved from the OSF database under accession code krv7g and 67tn3 (retrieved as OSF data sets osf.io/krv7g, and osf.io/67tn3).

## Code availability

The code accompanying the paper, specifying analyses and models is available on Github (111).

## Competing Interests

We have no conflict of interest to disclose.

## Acknowledgements

We are grateful to Megan McDonnell for helpful advice and discussion and assistance in data processing. We would like to thank the reviewers for helpful comments in improving the manuscript.

## Funding statement

We acknowledge grants PID2021-125453OB-I00 and PCI2025-167127-2 funded by MICIU/AEI /10.13039/501100011033 and co-financed by the European Union; grant AC20/00071 funded by Instituto de Salud Carlos III, CERCA Programme/Generalitat de Catalunya, and AGAUR/Generalitat de Catalunya (2017SGR01565, 2021SGR01522) (AC). MT is supported by MICIU/AEI (FPI program). Data collection for this work was supported by NSF NCS BCS 1734916/1954107 (to MAS and Byron Yu), NIH R01 MH118929 (MAS and BY), NIH R01 EB026953 (MAS, BY and Brent Doiron), NIH R01 EY022928 (MAS), and the Simons Foundation (BY). MAS was also supported by NIH R01 NS147766. This work was developed at the building Centro Esther Koplowitz, Barcelona.

## Author Contributions

Conceptualization: MT, MAS, JB, AC. Experimental Conceptualization: AU, RCW, MAS. Data curation: AU, RCW. Formal analysis: MT. Investigation: MT, JB, AC. Computational Methodology: MT, JB, AC. Experimental Methodology: AU, RCW, MAS. Software: MT. Visualization: MT. Writing–original draft: MT, JB, AC. Writing–review&editing: MT, MAS, JB, AC. Supervision: JB, AC. Funding: MAS, AC. All authors reviewed the paper for intellectual content.

## Notes

### Competing Interest Statement

The authors have declared no competing interest.

### Summary of Updates

Added Figure 7, evidence from a human multi-item working memory dataset supporting the model prediction of load-dependent storage strategy changes. Added Figure 6g, evidence from monkey behavioral and electrophysiological data supporting the prediction of increased precision associated with redundant storage. Updated Figure 3e, supplemental files. Minor changes to the text for clarity.

