## SupplementaryInformation for "Redundant prefrontal hemispheres adapt storage strategy to working memory demands"

Supplementary information

Melanie Tschiersch[1,2], Akash Umakantha[3,4] , Ryan C. Williamson[3,4], Matthew A. Smith[3,4,5], Joao Barbosa[6,7,8†] , Albert Compte[1,9†*]

[1]IDIBAPS, Barcelona, Spain,

[2]Programa de doctorat en Biomedicina, Universitat de Barcelona (UB), Barcelona, Spain,

[3]Center for the Neural Basis of Cognition, Pittsburgh PA, USA,

[4]Carnegie Mellon University Neuroscience Institute, Pittsburgh PA, USA,

[5]Carnegie Mellon University Department of Biomedical Engineering, Pittsburgh PA, USA,

[6]Laboratoire de Neurosciences Cognitives et Computationnelles, INSERM U960, Ecole Normale Superieure - PSL Research University, 75005, Paris, France,

[7]Cognitive Neuroimaging Unit, INSERM, CEA, CNRS, Université Paris-Saclay, NeuroSpin center, 91191 Gif/Yvette, France,

[8]Institut de neuromodulation, GHU Paris, psychiatrie et neurosciences, centre hospitalier Sainte-Anne, pôle hospitalo-universitaire 15, Université Paris Cité, Paris, France,

[9]Institut d'Investigacions Biomèdiques de Barcelona (IIBB), CSIC, Barcelona, Spain

[†] Equal contribution

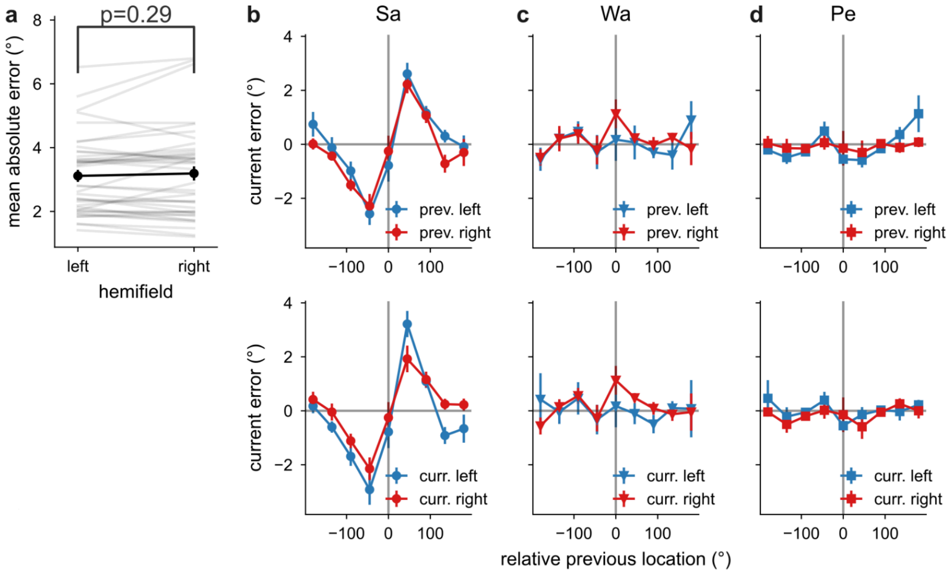


**Supplementary Fig. 1: Behavioral equivalence of left and right hemifield stimuli.** a) Mean absolute error for items presented in the left and right hemifield does not differ significantly (paired t-test: t(39) = -1.07, p = 0.29, mean = -0.08, 95% CI [-0.22,0.07]). b-d) Serial dependence curves for monkey Sa (b), Pe (c), Wa (d), split by whether the previous (top) or the current item (bottom) was presented in the left or the right hemifield. Tests were performed across N = 40 independent sessions of n = 3 monkeys. Error bars represent s.e.m.

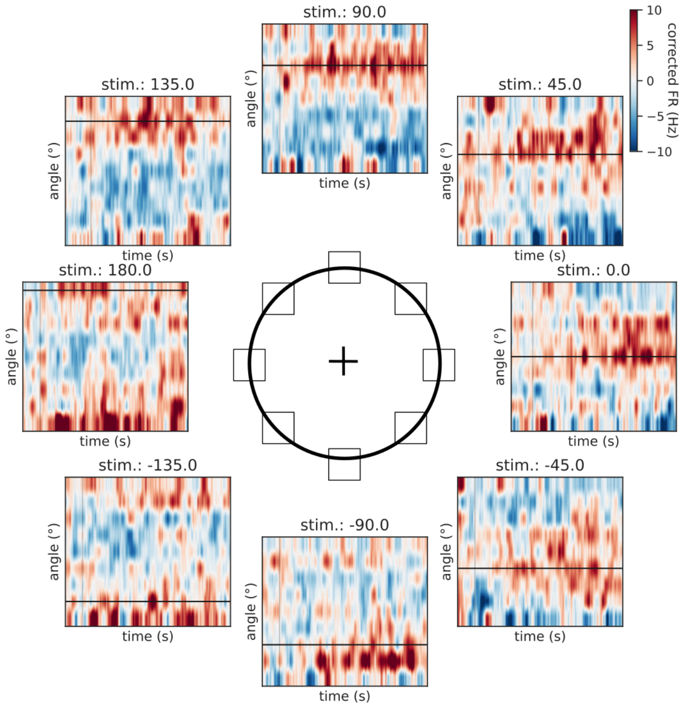


**Supplementary Fig. 2: Neural firing in example single trials with varying stimulus location.** Multi-units (y-axis) are grouped by their memory fields into 8 bins across circular space and the average of the multi-unit's mean-subtracted firing rates within each bin (win.: 200ms, step: 10ms) is shown over time. Colors are mapped for all targets in the same limits, see colorbar top right corner.

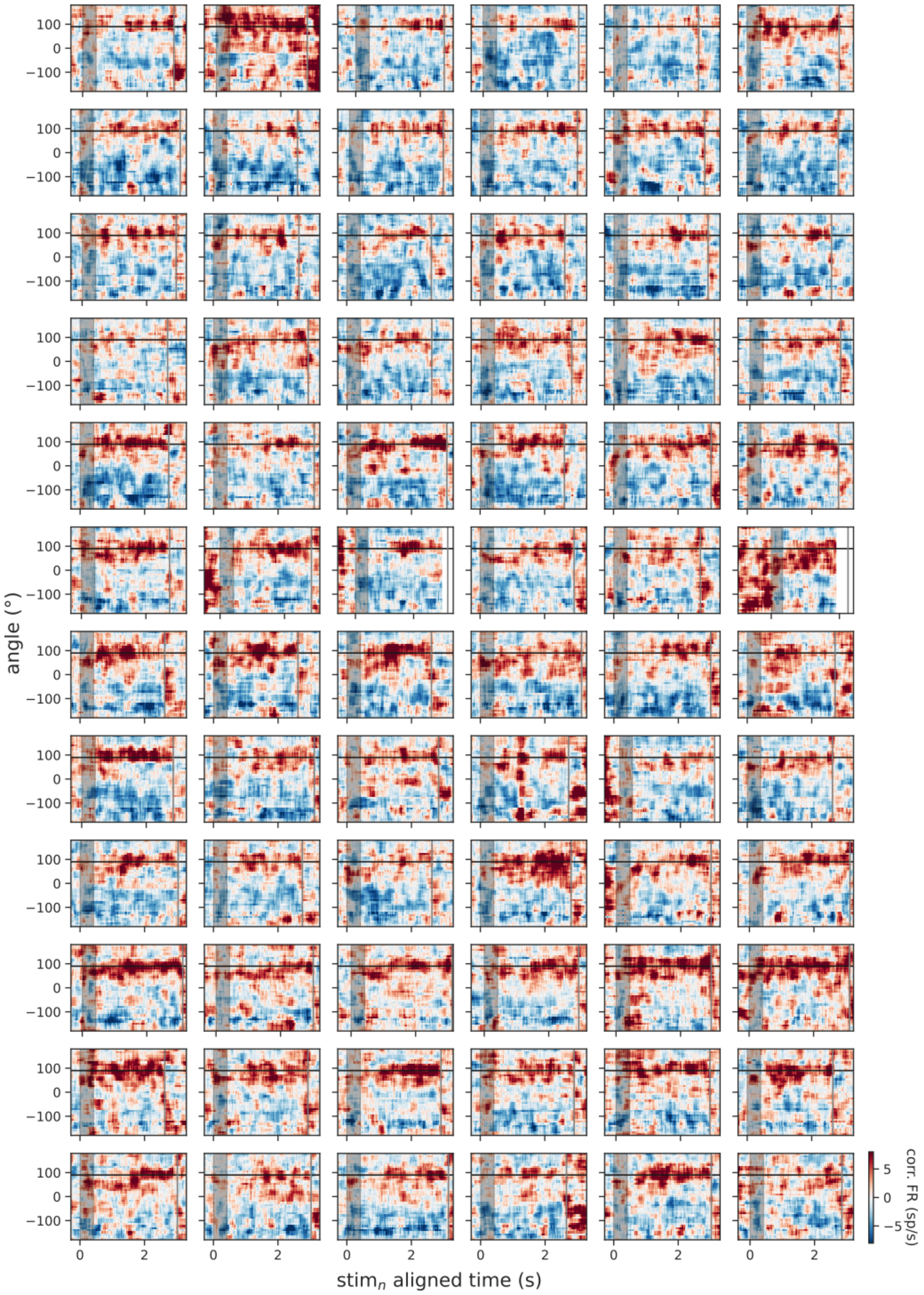


**Supplementary Fig. 3: Neural firing in all single trials with stimulus location at 90° in an example session (monkey: Sa, session: 0).**

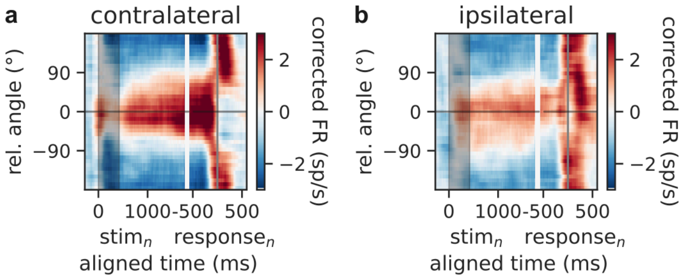


**Supplementary Fig. 4: Neural firing averaged across trials for a) contralateral and b) ipsilateral stimuli in an example session (monkey: Sa, session: 0).**

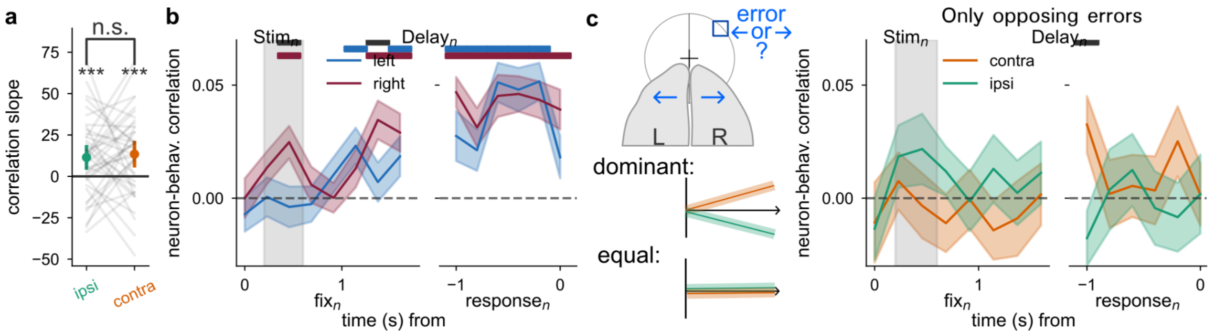


**Supplementary Fig. 5: Both hemispheres contribute equally to the response.** a) Quantification of the slopes of ipsilateral vs. contralateral correlation with behavioral response of Fig. 3b. One-sample two-sided t-test shows significance for ipsilateral: $t(39) = 3.07$, $p = 3.88 \times 10^{-3}$, 95% CI [3.91,19.01] and contralateral conditions: $t(39) = 3.34$, $p = 1.85 \times 10^{-3}$, 95% CI [5.3,21.55]. Paired two-sided t-test shows no significant difference between conditions $t(39) = 0.39$, $p = 0.7$, 95% CI [-8.34,12.27]. b) Full time course of the correlation of left vs. right hemisphere prediction errors with behavioral errors (slopes shown in Fig. 3c). c) Correlation of ipsilateral and contralateral prediction errors with behavior when hemispheres disagreed (e.g. counterclockwise (left hemisphere) vs clockwise (right hemisphere) prediction, left top). If a hemisphere would consistently dominate behavioral errors (here e.g. contralateral), behavioral-neural correlations should increase for the dominant side and decrease for the non-dominant side (left, middle). If hemispheres contribute equally, no such difference should emerge (left, bottom). When performing this analysis on the data, neither hemisphere seemed to dominate the behavioral correlations (right). Tests were performed across N = 40 independent sessions of n = 3 monkeys. Error bars are s.e.m., top bars indicate significance of time points (black: difference between left, right).

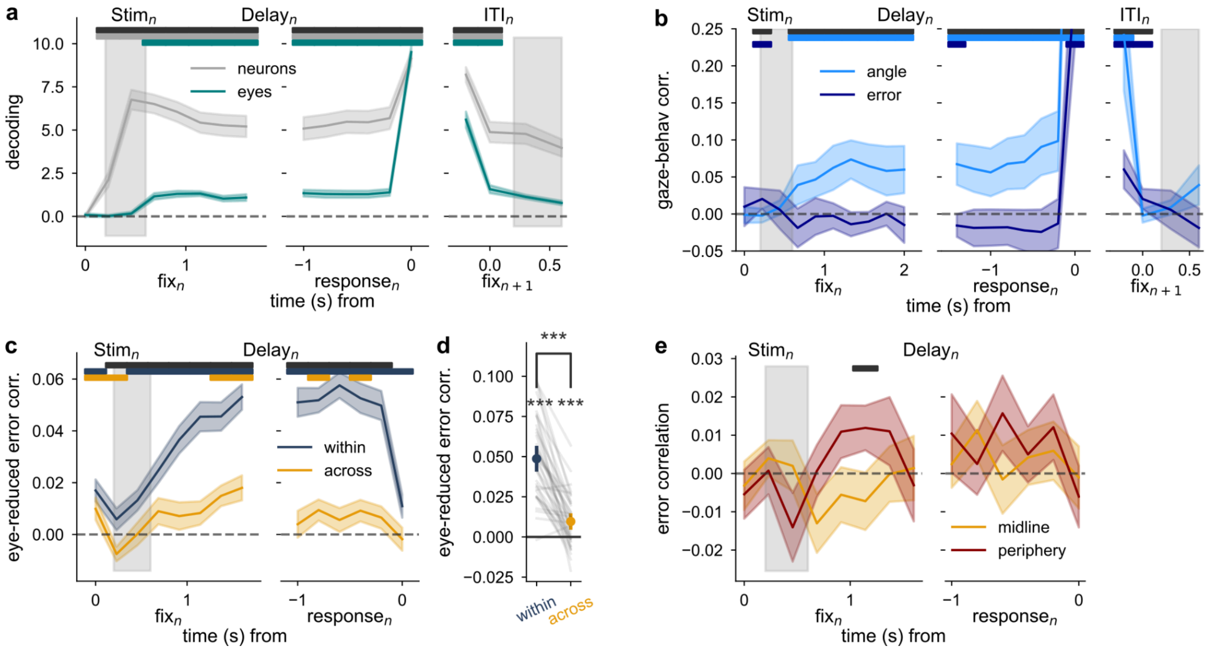


**Supplementary Fig. 6: Error correlations are not due to common input related to uninstructed eye movements.** a) A significant portion of the stimuli can be decoded from uninstructed eye movements even during the delay (while the monkey is fixating). However, this portion is smaller than the decoding from the multi-units. b) The angles of uninstructed eye movements are significantly correlated to target angles during the task. However, gaze errors are not correlated to behavioral errors during the delay, confirming that uninstructed eye movements do not provide common inputs informative about the errors correlated in Fig. 3e. c) Analysis from Fig. 3e but when introducing a regressor for eye movements during the training of the decoder. During testing only the angles inferred by the multi-units are interpreted. Error correlations across areas are still significantly above 0, excluding common input as an explanation. d) Quantification of average delay correlations (average of 7 independent 200ms bins) of panel c. One-sample two-sided t-test within: $t(38) = 12.11$, $p = 1.29\times10^{-14}$, 95% CI [0.04,0.06]; across: $t(38) = 3.76$, $p = 5.79\times10^{-4}$, 95% CI [0.0,0.01]. Paired two-sided t-test within to across: $t(38) = 9.73$, $p = 7.20\times10^{-12}$, 95% CI [0.03,0.05]. e) Across condition of Fig. 3e for trials presented on the vertical midline (90°, 270°) or in the periphery

(0°, 180°). Tests were performed across N = 39 independent sessions (1 session excluded due to shorter stimulus duration) of n = 3 monkeys. Error bars represent s.e.m. (panels a,c) or C.I. (panels (b, d).

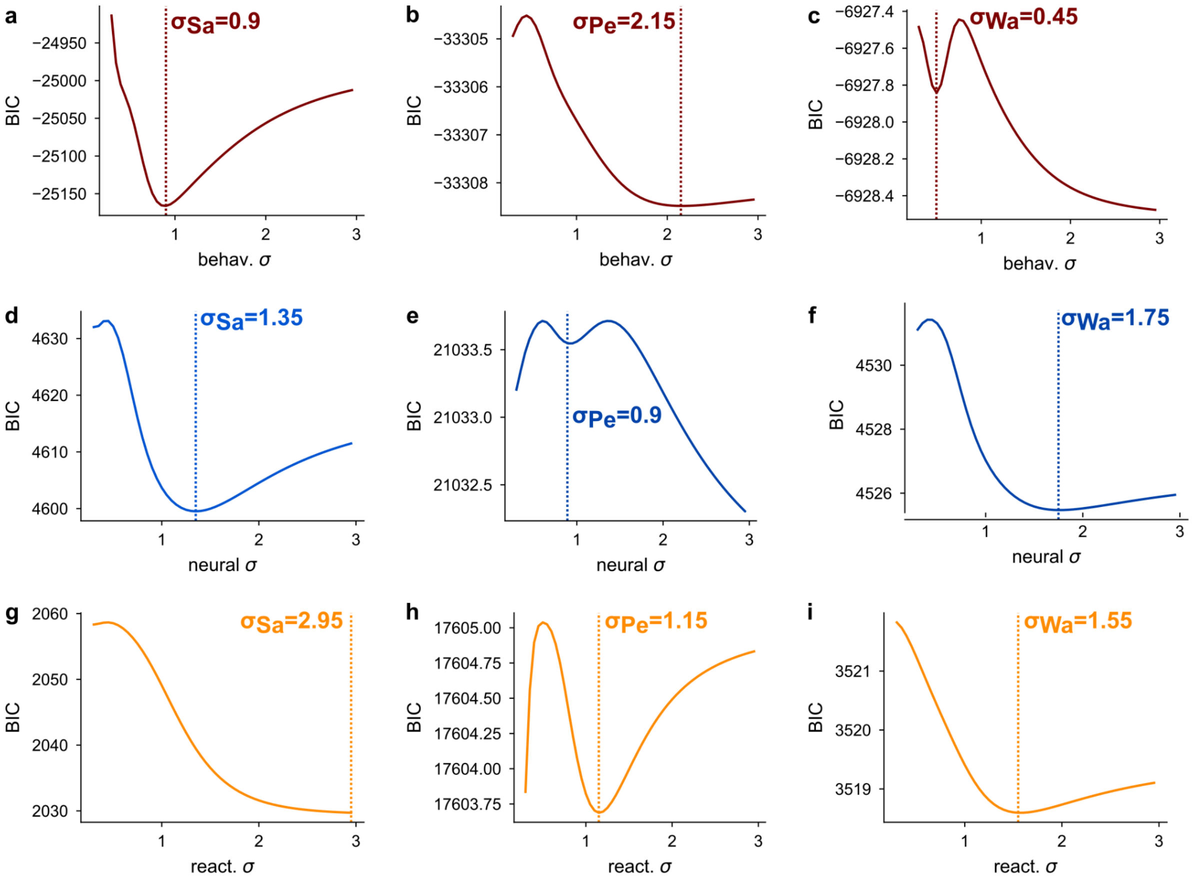


**Supplementary Fig. 7: BIC-score evaluation of best hyperparameter σ (width of the curve) for DoG1 model fits.** BIC score for different hyperparameters σ computed for a-c) behavioral, d-f) neural and g-i) reactivation attraction for each monkey (Sa: a,d,g; Pe: b,e,h; Wa: c,f,i). The best DoG1 fit is determined as the first occurring BIC minimum.

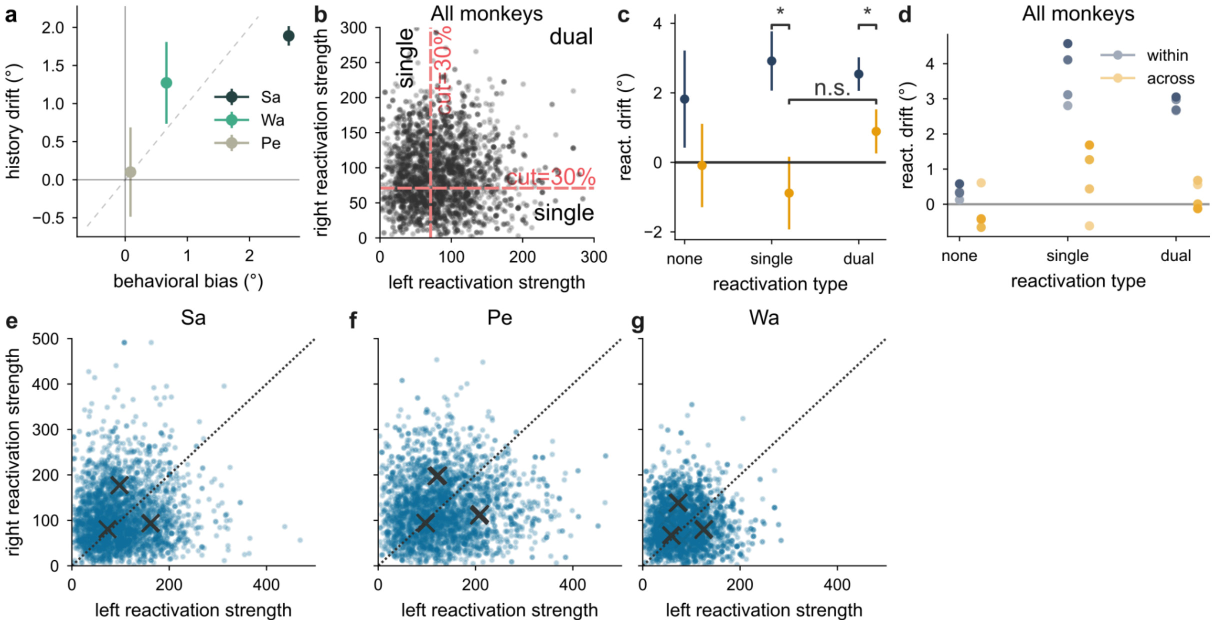


**Supplementary Fig. 8: Reactivations occur mostly independently in each hemisphere, as predicted by the model.** a) The strength of the neural history drift (extracted from decoder errors) covaries with the strength of the behavioral serial dependence across monkeys. b) Reactivation strengths extracted from the left, right hemisphere decoders shown for 1000 individual example trials. The cut-off of 30% of reactivation strength in each session is approximately indicated through red lines, splitting trials into high and low reactivation trials for each hemisphere. c) Average of last 400ms (2 independent 200ms bins) of reactivation drift shown in Fig. 5d when splitting trials based on their reactivation strength (see panel b) into null, single or dual reactivation trials (cut-off = 30% in each session). Reactivation drift only differed for within and across when at least one reactivation occurred (single, dual). Paired two-sided t-test None: $t(24) = 1.02$, $p = 0.31$, mean = 1.91, 95% CI [-1.95, 5.77], Single: $t(24) = 2.98$, $p = 0.004$, mean = 3.8, 95% CI [1.17, 6.43], Dual: $t(24) = 2.11$, $p = 0.04$, mean = 1.65, 95% CI [0.04, 3.26]. Across conditions between single and dual did not significantly differ from each other, suggesting that the dual class was produced by noise rather than a true difference in reactivation strengths. Paired two-sided t-test across dual-single: $t(24) = 1.33$, $p = 0.19$, mean = 3.42, 95% CI [1.05, 5.8]. Tests were performed across N = 23 independent sessions from n = 2 monkeys with positive behavioral serial dependence (monkeys Sa, Wa). d) Same as panel c but for different cut-offs. e,f,g)

Gaussian mixture model of fitting three classes on left vs. right reactivations. The classes were located around zero and off-diagonal, indicating that reactivations most often occur not at all or in only one of the hemispheres. Results are similar for the three monkeys Sa (e), Pe (f), (Wa) (g) and stable across different initial conditions (250 repetitions) for all three monkeys. Error bars represent s.e.m.

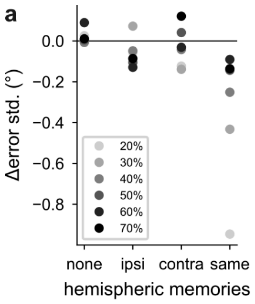


**Supplementary Fig. 9: Redundant hemispheres increase precision.** a) Different cut-offs for precise (absolute decoder error smaller than the cut-off) and imprecise memories in each hemisphere shows stability of results shown in Fig. 6g. Only averages are shown computed from N = 40 sessions of n = 3 monkeys.

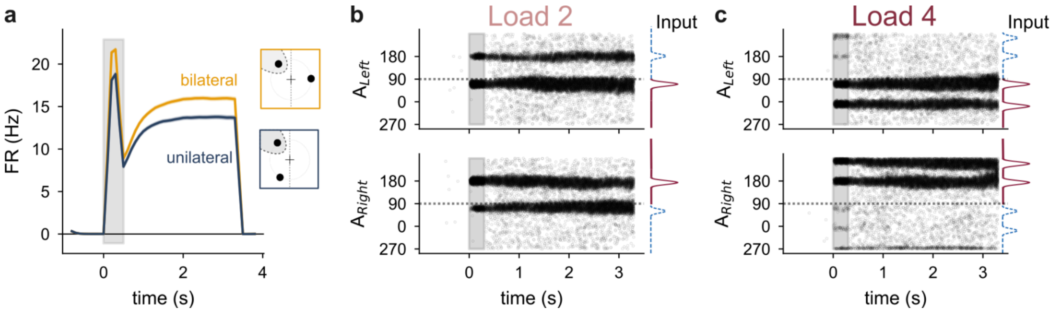


**Supplementary Fig. 10: High-capacity model simulations.** a) Average firing rate of neurons selective for a target location (window of 200 neurons around target location) in the load-2 condition is reduced when a non-target is presented in the target-hemifield (unilateral condition (n = 658 trials), see sketch on the right) in comparison to when a second item is added in the opposite hemifield (bilateral condition, n = 821 trials), akin to experimental results. b-c) Example simulations for a load-2 trial (b) and a load-4 trial (c) showcasing redundancy (both items stored in each area) and specialization (only contralateral items stored in each area).

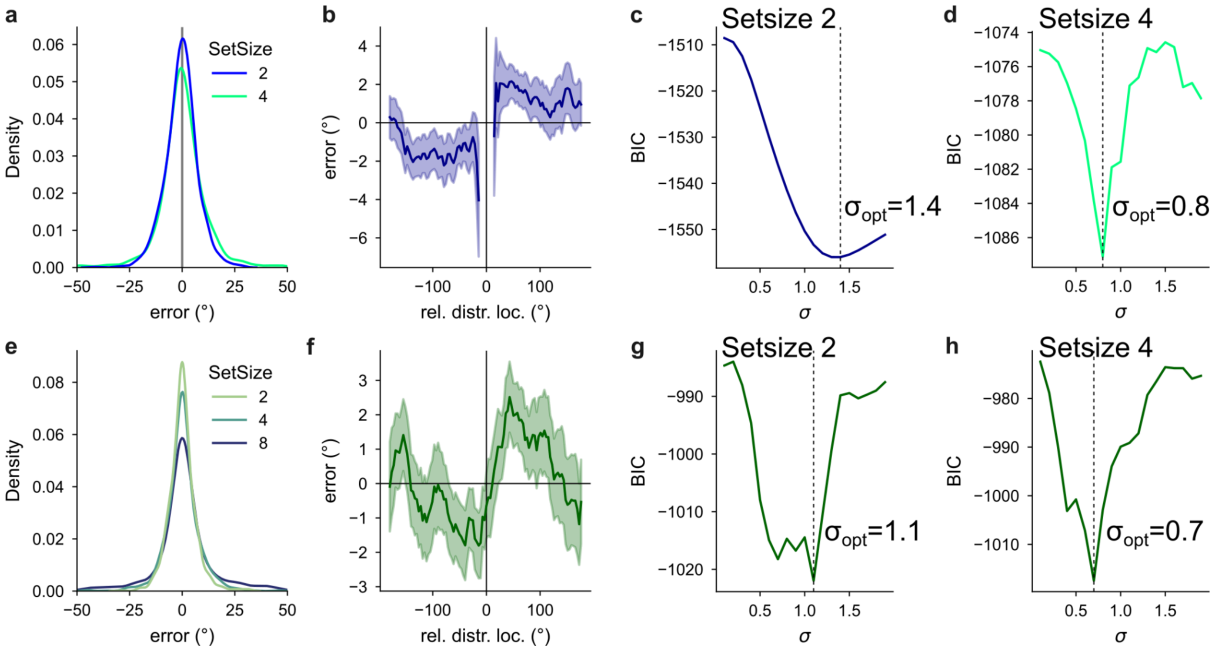


**Supplementary Fig. 11: Both human behavioral data sets are comparable.** a) Histogram of errors split by set size for Data1 (n = 10 subjects). b) Non-target attraction indicated by the error magnitude relative to the distance of the target and the strongest non-target for Data1. c) DoG fit to determine the strongest non-target and non-target attraction (see Methods) for load-2 trials. d) Same as panel c for load-4 trials. e-h) Same as panels a-d for Data2 (n = 8 subjects). Error bars are confidence intervals.
